# Deep sampling of ancestral genetic diversity reveals *Saccharomyces cerevisiae* pre-domestication life histories

**DOI:** 10.1101/2021.09.07.459046

**Authors:** Tracy J. Lee, Yu-Ching Liu, Wei-An Liu, Yu-Fei Lin, Hsin-Han Lee, Huei-Mien Ke, Jen-Pen Huang, Mei-Yeh Jade Lu, Chia-Lun Hsieh, Kuo-Fang Chung, Gianni Liti, Isheng Jason Tsai

## Abstract

The ecology and genetic diversity of model yeast *Saccharomyces cerevisiae* prior to human domestication remain poorly understood. Taiwan is regarded as part of this yeast’s geographic birthplace where the most divergent natural lineage was discovered. Here, we deep sampled the broad-leaf forests across this continental island to probe the ancestral species diversity. We found that *S. cerevisiae* is distributed ubiquitously at low abundance in the forests. Whole-genome sequencing of 121 isolates revealed nine distinct lineages, the highest known in any region. Three lineages are endemic to Taiwan and six are widespread in Asia. Molecular dating placed the divergence of the Taiwanese and Asian lineages during the Pleistocene, when a transient continental shelf land bridge connected Taiwan to other major landmasses. Extensive historical and recent admixture events were detected between natural lineages. In particular, the genetic component from a lineage associated with fruits that spanned the widest geographical range was present in most admixed isolates. Collectively, Taiwanese isolates harbor genetic diversity comparable to that of the whole Asia continent, and different lineages have coexisted at a fine spatial scale even on the same tree. Patterns of variations within each lineage revealed that *S. cerevisiae* is highly clonal and predominantly reproduces asexually in nature. We detected prevalent purifying selection genome-wide, with lineage-specific signals of positive or directional selection independent between lineages. This study establishes that *S. cerevisiae* has rich natural diversity sheltered from human influences, making it a powerful model system in microbial ecology.

## Introduction

The yeast genus *Saccharomyces*, which includes *S. cerevisiae*, is a powerful model system for revealing patterns of genomic variation underlying reproductive isolation and adaptation in eukaryotic microorganisms. Surveys of population genetic data have been used in *S. cerevisiae* to date the origin of key domestication events^1–3^, life cycle frequencies in nature^4^, the genomic basis of adaptation at continental scale^1,2^, and more recently to establish its geographical origin and dispersal history^5^. Phylogenomic analyses of the *Saccharomyces sensu stricto* complex and extensive sequencing of collections across the world suggest that *S. cerevisiae* originated in East Asia and underwent a single out-of-Asia event^2,3^. The 1,011 genome project—the most recent large scale yeast population genomic study—discovered that three wild isolates from Taiwan showed an unprecedented high genetic diversity compared to populations from the rest of the world^2^. Population genomics of 266 domestic and wild isolates in China revealed six wild lineages from primeval forests. The newly identified CHN-IX group represents the most diverged lineage^2^. Surprisingly, isolates from the CHN-IX group and the three Taiwanese isolates were in fact a single lineage that exhibited a disjunct geographic distribution^6^. Although considerable knowledge is available on the biogeography and population genetics of plants and animals across continents^7^, little is known about how eukaryotic microorganisms such as *S. cerevisiae* disperse, establish, reproduce and persist in nature^8^.

Most *S. cerevisiae* biology has been based on experiments on a handful of laboratory domesticated strains, but comprehensive analyses of the ecology and evolutionary biology of *S. cerevisiae* in the wild are still unavailable. In nature, *S. cerevisiae* have been isolated from the bark, fruits, surrounding soil, and leaves of plants belonging to several different families^9^, with early reports suggesting that the yeast is most successfully isolated from the oak Family Fagaceae^10–12^. *S. cerevisiae* contains high genetic diversity in certain populations, including lineage-specific variants that display clear population structures^1,2,10,13–19^ and explain phenotypic variance similar to common variants^20^. Samples from natural habitats tend to form unique populations with minimal genetic admixture, while lineages associated with human activities have either higher genetic admixture leading to a mosaic genome makeup or reduced genetic diversity after experiencing population bottlenecks^10,21–23^. The diverse natural lineages of *S. cerevisiae* present in East Asia provide an excellent opportunity to study the natural diversity of this species, which was previously believed to be fully domesticated^24^. Taiwan is a continental shelf island with the fifth highest tree density in the world^25^. Geological evidence indicates that the island underwent repeated reconnections to the Asian continent due to reduced sea levels during the Pliocene and Pleistocene glacial cycles^26^. The formation of land bridges thus allowed plants to readily migrate to Taiwan. Specifically, among the 13 climate-related forests types in Taiwan, five are Fagaceae-dominated natural forests on low- and mid-elevation mountains^27^, thus a potentially ideal natural habitat for *S. cerevisiae.* Taiwan also harbors a high phylogenetic diversity of flowering plants (53 out of 64 angiosperm orders present under the APG IV classification system^28^) and endemism compared to other oceanic islands^29^, raising the possibility that present lineages are genetically different from their continental counterparts.

Here, we set out to characterize the population-level diversity and distribution of *S. cerevisiae* in Taiwanese forests. We collected a total of 2,461 samples from a total of 1,101 plants, lichens, mushrooms and other materials at an elevational range from 0 to 3,118 meters above sea level. We used amplicon sequencing to characterize the relative abundances of *Saccharomyces* in a forest, and a total of 121 *S. cerevisiae* isolates were recovered with their whole genome sequenced. Combined with previously published studies, our study yielded a total of 137 genomes isolated from Taiwan. We provide evidence that genetically diverged lineages found in Asia are also present in Taiwan. These lineages have coexisted at a fine spatial scale, enabling us to study the pre-domestication phase of *S. cerevisiae* at an unprecedented resolution, including the extent of admixture between lineages. These results broaden our understanding of the ecological and biogeographic implications of a key model microorganism prior to anthropogenic impacts.

## Results

### Deep sampling of natural *S. cerevisiae* from Taiwanese forests

From July 2016 to October 2020, our sampling strategy consisted of maximizing the number of localities associated with Fagaceae hosts and sampling a broad range of plant families present in Taiwanese broad-leaved forests (**Fig. 1a**, **Supplementary Table 1**). We surveyed 693 plant hosts belonging to 43 orders, 86 families and 156 genera (**Supplementary Table 2**) collected over 113 non-overlapping 1 km^2^ grids. Various substrates (twigs, bark, leaves, flowers, fruits and topsoil around trees) were collected from each tree and subject to selective media enrichments resulting in 5,526 independent incubations (**Supplementary Table 3**). The successful isolation rates of *S. cerevisiae* per sample and per tree host was 1.9 and 10.8%, respectively, higher than from Brazilian forests, but lower than from North American oaks^12^ and Chinese wild niches^10^. These isolates were recovered across altitudes of 0–2,100 meters from 18 plant families (**Fig. 1b**), with a majority from Fagaceae including four genera (27 *Quercus*, nine *Lithocarpus*, eight *Castanopsis*, and one *Fagus* species). Interestingly, ten plant genera had higher isolation rates than *Quercus*, ranging from 40 to 100% per plant, albeit this recovery rate applied for as few as one tree (**Supplementary Table 2**). Among Fagaceae, *Quercus pachyloma* showed the highest isolation rate (75%; three out of four trees). Of the 339 lichen samples, four yielded successful isolations. Among the types of substrates, litter had the highest isolation rate (8.1%), providing the majority of recovered *S. cerevisiae* isolates (26.2%), followed by fruit, soil, bark and leaves (around 4–5% each). In general, the majority of samples were collected from July to December, and we found the isolation rate to be highest in July (18.9% per host tree), followed by September and October (17.5 and 11.3%, respectively). Isolation rates in other months remained around 0–11% (**Supplementary Table 3**).

**Fig. 1.**
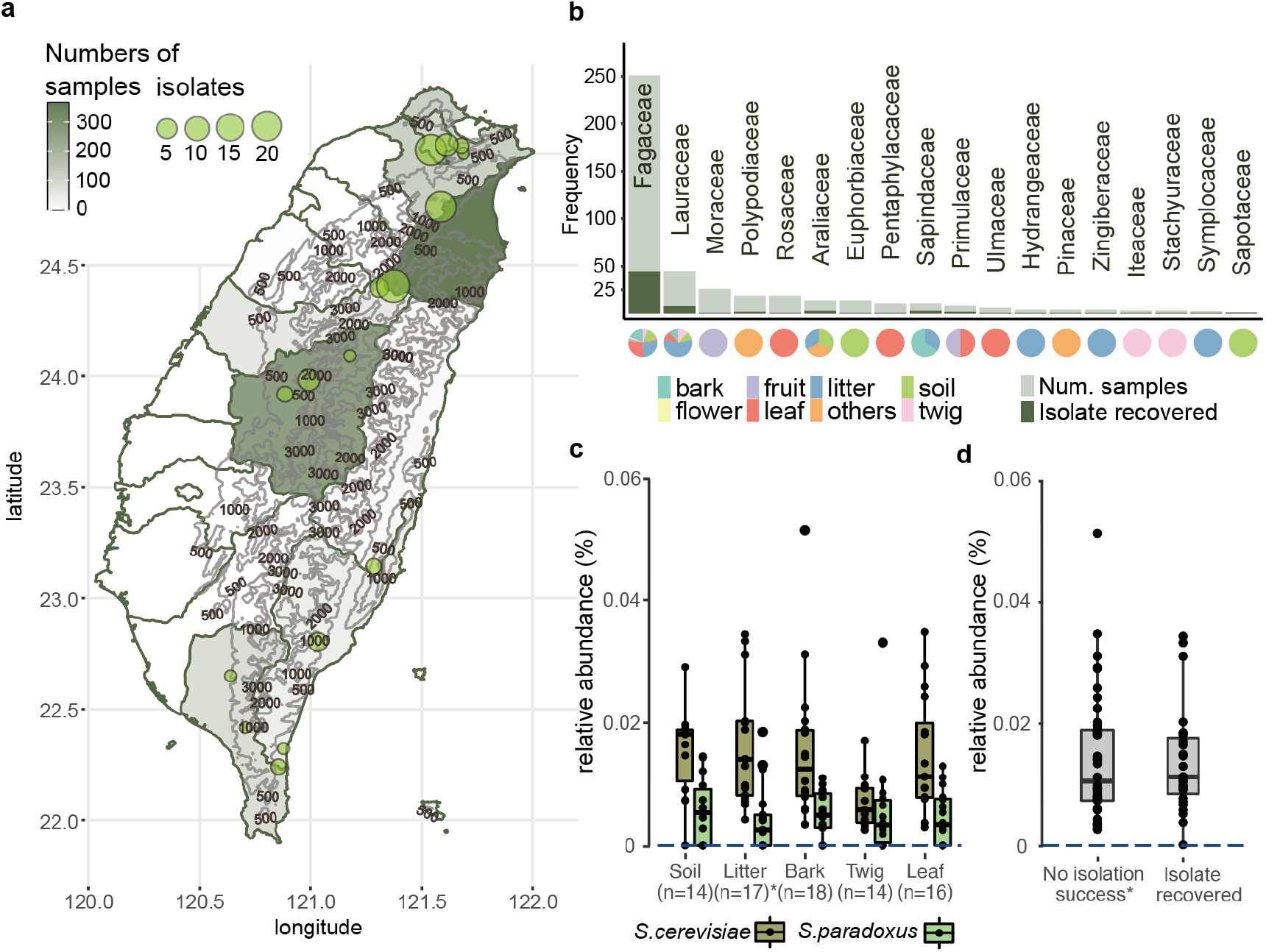
Deep sampling and isolation of *S. cerevisiae* in Taiwan. **a.** Map of Taiwan showing sampling efforts in each county, with darker shades representing areas with higher number of samples collected and circles denoting the locations where *S. cerevisiae* was successfully isolated. One isolate found on Dongsha Island is not shown on this map. **b.** Eighteen plant families from which *S. cerevisiae* was isolated. The darker color on each bar corresponds to the number of plants that yielded a successful isolation. Another 73 plant families from which we did not obtain any *S. cerevisiae* isolates are not shown. Pie charts below each bar represent the substrate surrounding plants from which samples were recovered. **c.** Pairwise comparisons found no differences in the relative abundances of *S. cerevisiae* among bark, leaf, twig (Wilcoxon-rank with Bonferroni correction, bark-leaf, P=1.0; bark-twig, P=0.118, leaf-twig, P=0.461) and **d.** between samples with or without isolation success.

One major challenge of characterizing the ecology of *Saccharomyces* is that isolations using selective enrichment media often do not accurately represent the yeast’s relative abundance in nature^30^. A repeated sampling of 8 trees over two years showed differential isolation successes (**Supplementary Table 4**), initially suggesting that *S. cerevisiae* had different abundances in different parts or trees. Focusing on a total of five substrates from 18 trees within ~100 m^2^ of this forest (**Supplementary Fig. 1 and Table 4**), ITS amplicon sequencing succeeded in detecting just two amplicon sequence variants (ASVs) belonging to the *Saccharomyces* genus—*S. cerevisiae* and *S. paradoxus.* In contrast to surveys in temperate and boreal forests^30–32^, *S. cerevisiae* had a higher relative abundance calculated as the percentage of the total taxa-classified reads than *S. paradoxus* in the subtropics (**Fig. 1c**). The sequence relative abundance of *S. cerevisiae* was on average 0.012% in these trees belonging to seven families regardless of substrates sampled; this suggested that, despite being ubiquitous in nature, *S. cerevisiae* lives in small populations. The relative abundances of *S. cerevisiae* were found to be constant between pairwise comparisons of bark, leaves and twigs (**Fig. 1c**; Wilcoxon-rank with Bonferroni correction, bark-leaf P=1.0; bark-twig P=0.118; leaf-twig P=0.461); among tree families (**Supplementary Fig. 2,** P=1.0); and whether a *S. cerevisiae* isolate was recovered (**Fig. 1d;** P=0.89). In addition, bioclimatic variables extracted from GPS coordinates also exhibited no difference between sites at which isolates were and were not recovered (**Supplementary Info and Supplementary Table 5**). Together, these results imply that the primary habitat of *S. cerevisiae* is unlikely associated with a single tree host.

### Multiple natural *S. cerevisiae* lineages in Taiwan

We sequenced the genomes of 121 isolates with a median coverage of 91X depth (**Supplementary Table 6**). All isolates were primarily homozygous (average heterozygosity: 0.01%) diploids, with the exception of isolate PD36A, which was a triploid (**Supplementary Fig. 3**) estimated by flow cytometry (**Supplementary Info**). We constructed a maximum likelihood phylogeny based on 808,864 SNPs segregating in 340 isolates (**Fig. 2a**) by including of 219 representative isolates previously studied from multiple habitats^1,2,33,34^ that sampled all the major world-wide wild and domesticated lineages. The topology of the isolate phylogeny is largely consistent with a previous neighbor joining tree from the 1,011 *S. cerevisiae* genome project^1^: the natural isolates were mostly grouped according to sampling locations, while industrial isolates were grouped according to fermentation sources. In particular, the Wine/European clade and Asian Fermentation clade were separated by a suite of natural isolates, suggesting independent domestication events^15,22,35^. The African Palm Wine clade was separated from the West African Cocoa clade and placed near the branch leading to the Asian Fermentation clade. Furthermore, the CHN-VI/VII clade, which was collected from fruits, was further separated into two clades consistently with geographical proximity of its members (designated as CHN-VI/VII.1 and CHN-VI/VII.2 in **Fig. 2a and 2c, Supplementary Table 6**).

**Fig. 2.**
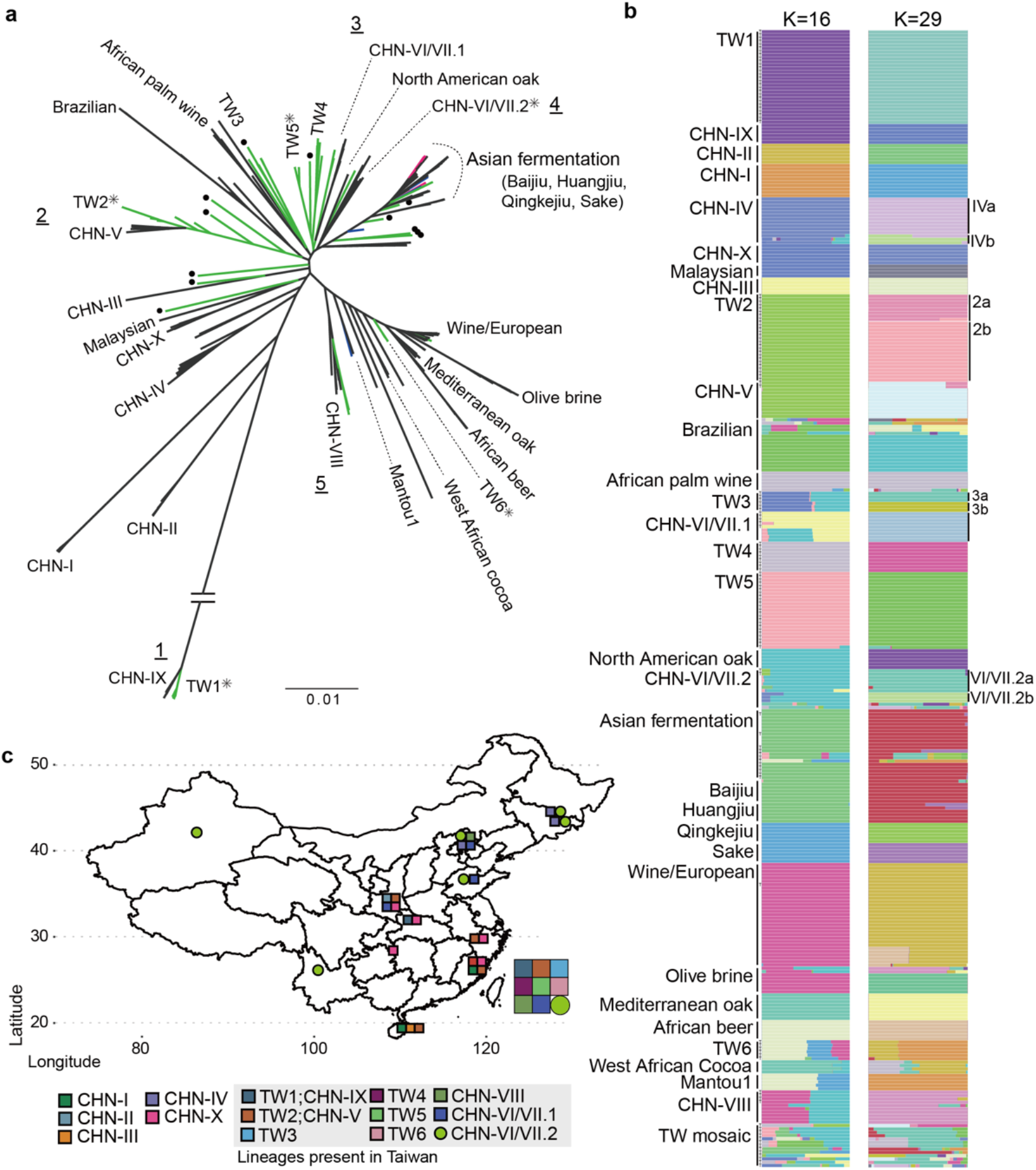
Phylogeny and population structures of 340 *S. cerevisiae* isolates. **a.** Unrooted phylogeny based on 808,864 genome-wide SNPs. Bootstrap support was >90% in all major clades except inner nodes within some clades indicated by asterisks. Natural, industrial and fermentation-related isolates discovered in Taiwan are in colored in green, blue and magenta, respectively. Five cases in which Taiwanese and Chinese isolates were found to be monophyletic are indicated with underscored numbers. The Asian fermentation clade includes Baijiu-, Huangjiu-, Qingkejiu-, Sake- and fermentation-related isolates from Taiwan, as shown in **b**. Mosaic Taiwanese isolates from ADMIXTURE analyses are labelled with circles on branch tips. **b.** Population structure from ADMIXTURE analysis at K=16 and 29. Labels on the left side of the bars indicate each group from K=16 and some were further separated in K=29 which are annotated on the right side. Natural Taiwanese isolates with admixed genome makeup are shown together in the TW mosaic group. **c.** Map of China and Taiwan indicating sampling locations of *S. cerevisiae* colored according to natural lineages. CHN-IV isolates that were sampled from Japan are not shown on this map.

Previous studies of natural *S. cerevisiae* revealed that most clades comprise isolates from neighboring geographic origins^1,2^; however, natural Taiwanese isolates are found throughout the phylogeny despite the small size of the island (**Fig. 2a**). The population structure of the 340 isolates used for the phylogeny were analyzed using ADMIXTURE^36^ with K from 2 to 30. Cross validation (CV) error was lowest at K=29 (CV error = 0.09025), though it only differed <1% between K=16 and 30 (**Fig. 2b, Supplementary Fig. 4**). ADMIXTURE at K=16 was largely consistent with the phylogenetic clades such as placing CHN-VI/VII into two genetic groups. ADMIXTURE at K=29 further separated two instances in which a group was split into solely either Chinese or Taiwanese isolates, suggesting the presence of lineage-specific segregating sites as a result of geographical isolation (**Fig. 2b and Supplementary Table 7**). Some groups comprising isolates from a proximate geographical origin were further split into smaller groups, suggesting ongoing genetic differentiation. Based on ADMIXTURE K=29, we reused previously assigned group names^1,2^ and designated these differentiated groups as well as new lineages exclusively found in Taiwan TW1 to TW6 (**Fig. 2**). Examples include the recovery of 28 TW1 isolates clustered with CHN-IX^2,6^, together representing the most divergent lineage to date, and a new TW4 group that did not contain any Chinese strains (**Fig. 2**). Interestingly, TW4 included isolates sampled from lichens and four isolates sampled from mushrooms that were previously placed in an undefined clade^1^, suggesting a possible symbiotic relationships with other fungi^37^. In other instances, Taiwanese isolates were found in three previously assigned groups such as CHN-VI/VII.1, CHN-VI/VII.2 and CHN-VIII. Isolates of the most diverged TW1/CHN-IX lineage were separated by approximately 1400 km, with four other natural lineages (CHN-I, V, VI/VII, and X) in between. Twenty-three isolates from northern Taiwan (TW2) were clustered with the CHN-V population sampled as far as 1500 km apart. Together, these results suggest that Taiwan harbors the highest number of lineages that exhibit disjunct distributions followed by the Hubei-Shanxi region (nine and five, respectively; **Fig. 2c**)

### Extensive admixture of natural lineages

Both inter- and intra-species spontaneous hybridizations have been documented in *Saccharomyces* species. For instance, the wild *S. paradoxus* SpC* lineage present in North America^38^ and domesticated *S. cerevisiae* Alpechin lineage^39^ are classic examples of past hybridizations that played genomic and phenotypic diversities^1,2,33,38^. Most Taiwanese isolates tend to have little admixture, with 20 and 5% (27/137, 7/137) of isolates containing at least 10% of the genetic component from two and at least three genetic ancestries (**Fig. 2b, Supplementary Table 7**), respectively. We confirmed the genetic components of domesticated strains’ origins in wild isolates from African cocoa^1^, olive brines and Brazilian forests^33^, and identified an additional TW4 group sharing major genetic components with the steamed buns (Mantou) and Wine/European clades, albeit recovered from nature. Other Taiwanese admixed isolates were apparent on the phylogenetic tree as isolated branches and had different levels of admixture from domesticated clades (**Fig. 2a**). Interestingly, 84% of admixed isolates harbor over 10% of their genetic components from CHNVI/VII-2a. Isolates containing admixed genetic components were placed in the phylogeny between the domesticated and wild clades such as the North American Oaks, TW5 and CHNVIII group. In addition, all Taiwan isolates recovered from fruits contain CHNVI/VII-2a genetic component (**Supplementary Fig. 5**). We further visualized the effective migration patterns of the Asian isolates using EEMS^40^, which inferred a region of high effective migration where isolates of CHNVI/VII-2a were present. Together with the non-admixed CHNVI/VII-2a isolates spanned the widest geographically distribution (**Fig. 2c**) suggested that fruits were agents of long-distance passive dispersal (**Supplementary Fig. 6**).

To confirm that gene flow occurred between genetic groups, we applied TreeMix^41^ to designated groups from ADMIXTURE K=16 (**Fig. 3a, Supplementary Info and Fig. 7**). The TreeMix phylogeny first indicated extensive gene flow among domesticated lineages such as solid- and liquid- state fermentation products and between natural lineages sister to domesticated lineages. Examples include isolates from steamed buns (Mantou) and Asian alcoholic beverages (Sake and Qingkejiu), as well as TW6 forest isolates. Second, the phylogeny also identified gene flow between natural lineages sister to the Wine/European and Asian Fermentation clades. The CHN-VIII group emerged from both the Wine/European and fruit enriched CHN-VI/VII-2 lineages, which contain isolates from fruits and the natural environment across the Asian continent, including Taiwan. Strikingly, we also recovered hybrids between natural lineages that coexisted in proximity. A striking example is the isolate PD38A, recovered from *Castanopsis fargesii* fruits (**Fig. 3a**). Two isolates each belonging to TW4 or TW2 clade came from fallen fruit, while PD38A was isolated from fruit growing on the tree. This PD38A hybridization timing was likely to be recent, given the presence of large haplotype blocks not extensively broken down by recombination containing variants identical to each parental lineage (**Supplementary Fig. 8**). Overall, these results suggest that hybridizations were common in *S. cerevisiae* and that some admixed lineages have persisted in nature. Reanalysis of the TreeMix phylogeny based on ADMIXTURE group K=29 shows consistent results—recurrent migrations occurred between lineages, leading to the Wine/European and Asian Fermentation clades (**Supplementary Info, Supplementary Fig. 9 and 10**).

**Fig. 3.**
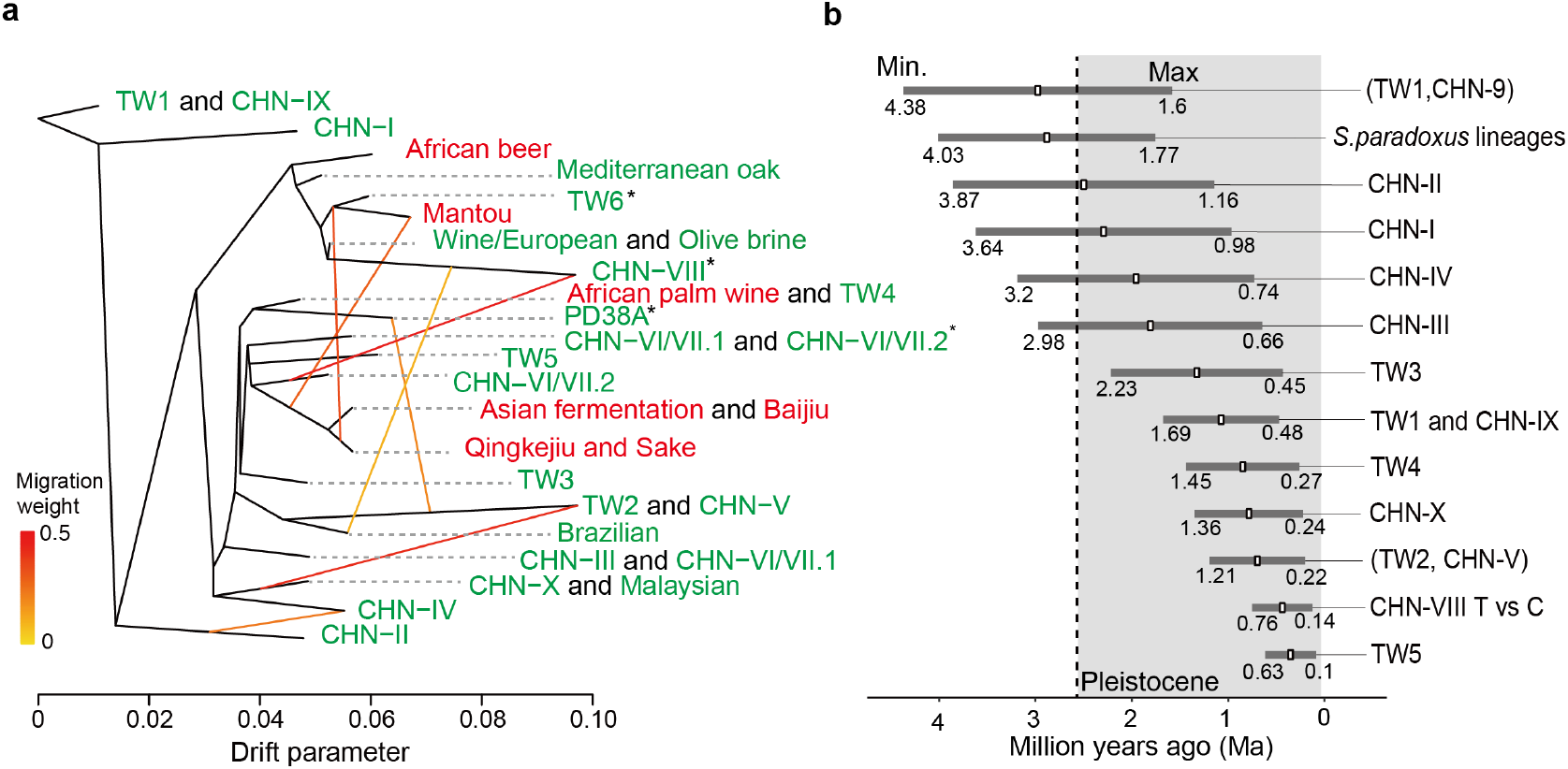
Migration and divergence time between lineages. **a.** Migration edges (yellow to red colored lines) estimated by TreeMix showing seven migration edges on the phylogeny. Different edge colors indicate the strength of migration. Lineages were colored according to isolation sources (red and blue denote domesticated and natural environment, respectively). Asterisk denotes lineages that contain multiple genetic components from different K from the ADMIXTURE analyses. **b.** Molecular estimate of time to the most recent common ancestor (Ma, million years ago) in different *S. cerevisiae* lineages. The full phylogeny is shown in **Supplementary Fig. 11**.

To incorporate these findings into a comparative resource, we further sequenced the genomes of 24 Taiwanese isolates using Oxford Nanopore reads (**Supplementary Table 8**). These genomes were chosen because they represent all the natural lineages discovered in Taiwan and five admixed isolates found in this study, including the recent natural hybrid PD38A. These genomes allowed us to estimate the divergence times of different lineages and the Chinese/Taiwanese split in greater detail; this is important because inferring population history in *S. cerevisiae* with different frequencies of asexual and sexual generations^4^ is challenging when using population genetics methods designed around human heterozygosity and recombination rates^42^. A phylogenetic tree of 62 *S. cerevisiae* isolates using five species of *Saccharomyces sensu stricto* as outgroups was recovered through coalescence-based analysis using the 1,594 single copy gene trees (**Supplementary Fig. 11**). The tree’s topology was consistent with our phylogeny inferred from genome-wide concatenated SNPs and the Chinese/Taiwanese splits remained robust with higher bootstrap support. Using MCMCtree with four molecular calibrations, we found the divergence of different natural lineages to be 0.1–4.38 million years ago (Ma) (**Fig. 3b; Supplementary Table 9**). In particular, we were interested in the age of the most recent common ancestor of TW1 and CHN-IX, which was estimated to have diverged from the rest of *S. cerevisiae* 1.59–4.38 Ma, then split 0.48–1.69 Ma during the Pleistocene epoch (**Fig. 3b**). The estimated divergence date at which the Taiwanese and Chinese isolates were unambiguously separated such as CHN-VIII lineage was also congruent with this estimate (0.14-0.86 Ma, **Fig. 3b**), suggesting that the split may represent a vicariant event resulting from the submergence of the Taiwan Strait Land Bridge during interglacial periods and/or uplift of Taiwanese mountains^43^ during this period.

### Biogeography and life history of wild *S. cerevisiae* lineages

To further assess the spatial relationships of isolates between and within the natural lineages, we closely examined the genetic diversity and range distributions of the Taiwanese isolates. Previous studies have shown that isolates within a lineage are not necessarily geographically proximate; on the other hand, secondary contact was observed in North America, where members of *Saccharomyces paradoxus* group A were sympatric with group B in one tree^44^. We identified multiple Taiwanese *S. cerevisiae* lineages co-occurred at the same locality (**Fig. 4a**, **Supplementary Figure 12 and Table 6**) even on the same tree. In one sampling area, four lineages were recovered less than 35 km apart in central Taiwan (TW1-TW4 and mosaics, n=10; **Supplementary Fig. 12**). In another sampling site—Fushan Botanical Garden, where we obtained 23 isolates comprising three lineages and admixed isolates were recovered (**Fig. 4a**). Both significant negative and positive correlation between genetic and geographical distance were observed in isolate pairwise comparisons in close distances (P<0.05 with 1,000 permutations; **Supplementary Fig. 13**). However, no such association was found of the whole region (**Fig. 4b**; Mantel’s r = 0.07, P=0.23), suggesting that in a given region the relationships between isolates was less determined by population structure of single lineages but dictated by the heterogeneity of multiple lineages coexisted at small spatial scale. Interestingly, the admixed isolates did not contain genetic components from adjacent isolates, but instead from CHN-VI/VIII.2a and others (**Supplementary Fig. 14**). In addition, these combinations of coexisting lineages were not present in a similar locality range in China (**Fig. 2c**), suggesting coexisting of lineages was established by independent dispersal events.

**Fig. 4.**
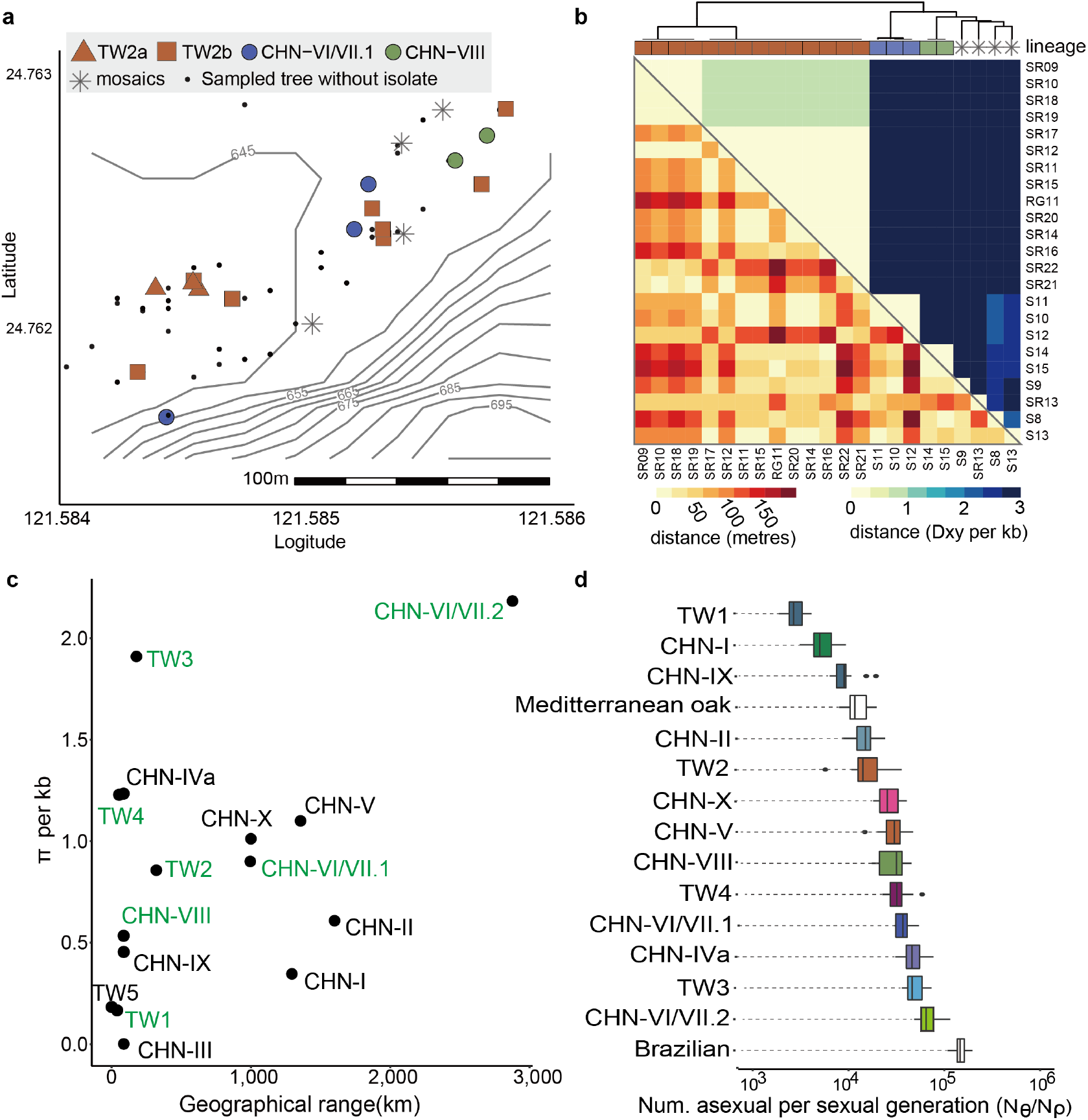
Patterns of genetic variations and geographical distribution in *S. cerevisiae*. **a.** Fine-scale geographic sampling at Fushan Botanical Garden in Taiwan. A total of 106 tree sites constituting 286 substrates were sampled in this region. Different colors represent different lineages and filled circle denote sampled trees from which *S. cerevisiae* was not successfully isolated. **b.** Genetic and geographic distance of isolate pairs identified in **a**. **c.** Lack of correlation between genetic differentiation θ_π_ and geographical range across lineages. Clones were excluded from this analysis. **d.** Frequency of asexual per sexual generations across lineages. Lineage TW5, TW6, CHN-III and North American oak were excluded due to either insufficient non-clonal individuals within a lineage or the majority of variations being non-informative singletons.

Our finding that divergent lineages occur in proximity implies that *Saccharomyces cerevisiae* exhibits a non-panmictic population structure that deviates from the standard neutral model. The overall genetic diversity of Taiwanese isolates was comparable to that of Chinese isolates (Taiwan θ_π_=5×10^−3^ versus China θ_π_=6×10^−3^), even though the samples were only meters to tens of kilometers apart (**Supplementary Fig. 15**). This reinforced that the pattern of *S. cerevisiae* diversity in a geographical region was shaped by the presence of multiple lineages and heterogeneity of metapopulations in the same habitat. In contrast, *S. cerevisiae* lineages were previously determined to be reproductively isolated from each other as a result of pre- and post-zygotic barriers^10,45^, and hence may more accurately determine different population genetics parameters. Up to a two-fold difference was observed in genetic diversity between lineages, with the aforementioned most widespread CHN-VI/VII.2a group harboring the greatest diversity (**Fig. 4c; Supplementary Table 10**). In contrast, when comparing isolates on the same tree at an extreme microgeographic scale, we found instances of all isolates being clonal or from different lineages with pairwise differences differed by ~35,000-fold (1–35,922 maximum number of pairwise mismatches of isolates recovered on the same tree; θ_π_=8.3×10^−8^–2.9×10^−3^; **Supplementary Table 11**). Three out of seven lineages have exhibited a linear isolation-by-distance (IBD)^46^ signature including the aforementioned TW2 lineage (p<0.05; **Supplementary Fig. 16**). TW2 lineage exhibited a central-southern Taiwan discontinuous distribution, where isolates are found as much as 194 km apart. This suggests that the greater the geographical range, the higher the likelihood of genetic differentiation. Indeed, greater sequence divergence was shown when intra-lineage isolates between lineages were >10 km apart (P<0.001; Wilcoxon rank-sum test; **Supplementary Fig. 17**), which supported genetic differentiation as a result of geographical isolation^47^.

Patterns of segregating sites can be used to infer the relative contributions and frequencies of reproduction modes in nature^4^. Wild *S. cerevisiae* isolates were highly inbred: Wright’s inbreeding coefficient *F* was an average of 0.99 and clones made up 16–100% of each lineage (**Supplementary Table 6**), suggesting that most generations were mitotic regardless of lineage. We estimated that the effective population size (N_e_) of mutational and recombination diversity for all chromosomes was 292,748–4,879,320 and 33–106, respectively, across lineages (**Supplementary Table 12**). The differences between both Ne estimates equates to approximately 2×10^3^–1×10^6^ mitotic cell division occurred for every meiosis (**Fig. 4d**). Such estimates overlap with previous estimates of 12,500–62,500 clonal generations based on the decay of heterozygosity during mitosis^48^, 1,000–3,000 in two genealogically independent populations of *S. paradoxus*^4^ and 800,000 generations in the fission yeast *Schizosaccharomyces pombe*^49^.

### Prevalent negative selection across lineages

The presence of different levels of shared genetic components observed between Chinese and Taiwanese isolates among the five shared lineages suggested a distinct differentiation between the disjunct populations. The average ratio of nonsynonymous to synonymous substitution rates *(dN/dS)* between China and Taiwan isolates across lineages was 0.21 (**Fig. 5a**), suggesting that there was prevalent negative selection acting on the proteome of *S. cerevisiae*, with only 40–303 out of 6,572 genes showing signals of positive or balancing selection (*dN/dS* > 1) across the Taiwanese lineages. Most of these genes were lineage-specific, with only *AIM21*, involved in mitochondrial inheritance, detected in four out of five lineages (**Supplementary Fig. 18 and Table 13**) suggesting that selection acted independently in these lineages. We next identified 89–3,677 highly differentiated SNPs (F_ST_=1) between Chinese and Taiwanese isolates across lineages (**Supplementary Table 14**), suggesting different levels of ongoing differentiation. These fixed sites with high F_ST_ among the four groups generally occurred at a low allele frequency (7–18%, **Supplementary Fig. 19**) in other lineages and harbored a low proportion of nonsynonymous variants (5–29%). Principle component analysis (PCA) of the frequencies in the lineage-differentiated alleles showed that the Chinese and Taiwanese isolates were clearly separated from each other and other lineages, suggesting that the direction of fixation was region independent (**Fig. 5b**). Together, these results show that a large fractions of genetic diversity present in the Taiwanese isolates reflect ancestral alleles shared with Chinese isolates were maintained by purifying selection and new genetic variants since divergence emerged independently across lineages despite coexisting in Taiwan.

**Fig. 5.**
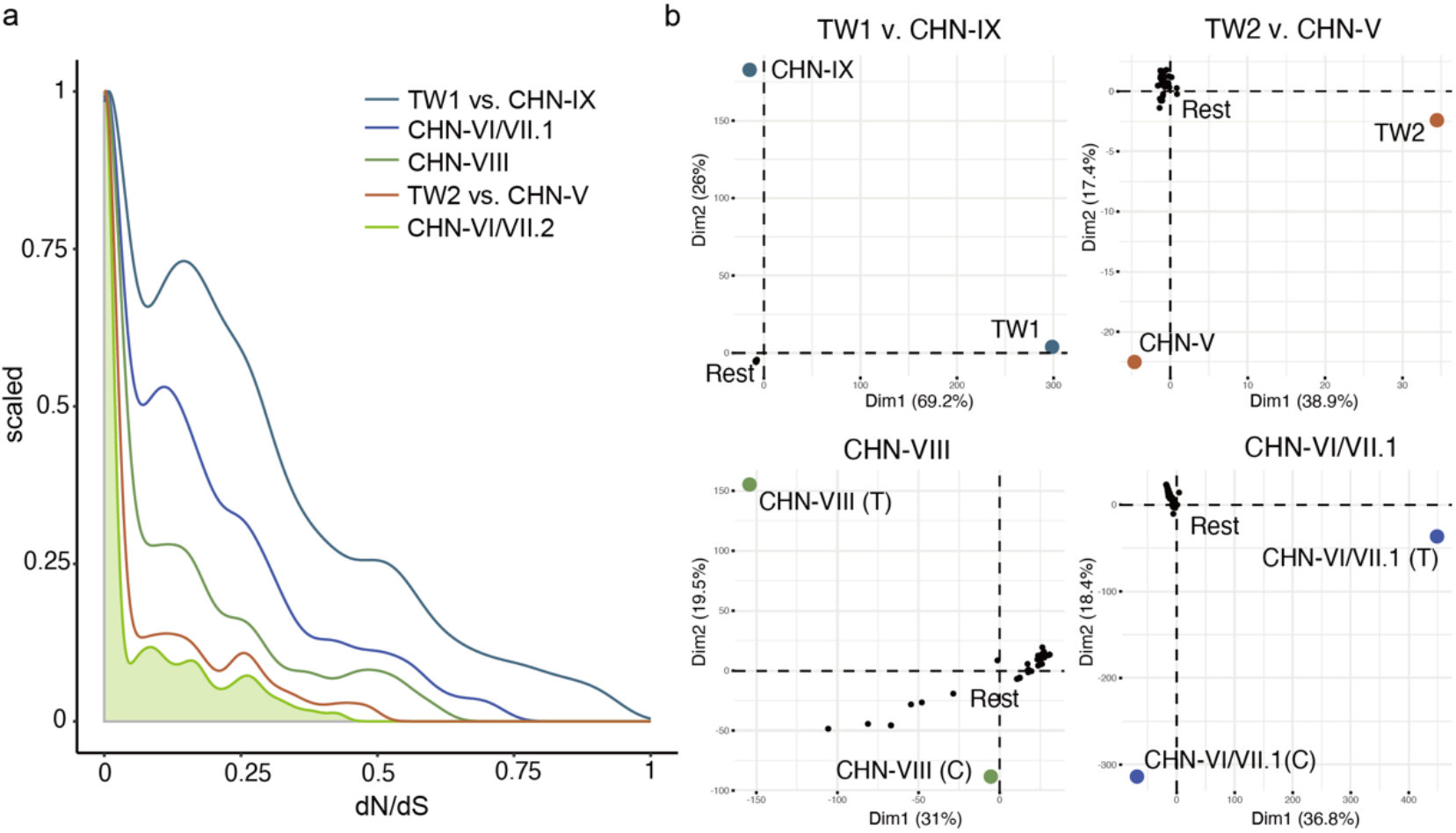
Prevalence of negative selection across lineages. **a.** Density plot of *d_N_/d_S_* showing the majority of genes with *dN/dS* < 1. **b.** Principal component analysis of allele frequencies in the highly differentiated (FST = 1) SNPs between Chinese and Taiwanese isolates in each lineage were clearly separate from each other and the rest of the lineages. Lineage CHN-VI/VII.2 was excluded since allele frequencies were unavailable because only one unique isolate was found in Taiwan.

## Discussion

A comprehensive understanding of the natural history of the budding yeast *S. cerevisiae* is key to further utilizing one of the most human-exploited microorganisms. In this study, we leveraged a four-year deep sampling in Taiwan and combined metabarcoding approach to uncover *S. cerevisiae*’s ubiquitous presence but low abundance in broadleaf forests. We isolated and whole-genome sequenced 121 isolates to confirm the presence of the most diverged lineage TW1^6^ and uncover five additional lineages that shared ancestries with lineages found in China as well as four new lineages exclusively found in Taiwan. We show that sympatric lineages coexist in different parts of Taiwan and identified frequent introgressions between lineages. We found that the population structure of *S. cerevisiae* can be explained by a markup of different lineages that each outcrossed on average once in every 3.0×10^3^–1.5×10^5^ meiotic generations. Despite living a predominantly asexual lifestyle, the species has a large effective population size, which was inferred to be the result of efficient purifying selection purging deleterious alleles^50^. The availability of high-quality *S. cerevisiae* assemblies presented here, in addition to genetic, molecular tools and genome resources such as the 1,011 genome collection^3^ already available in this model organism provides an exciting new platform to study microbial ecology.

Although *S. cerevisiae* has repeatedly been recovered from oak bark in the Northern Hemisphere^12,51^ and being the only substrate of isolation in recent studies^2,52^, our findings confirm that *S. cerevisiae* at the species level is consistent with the first part of Baas-Becking’s hypothesis: “everything is everywhere.” In addition to temperature, we speculate that isolation success for *S. cerevisiae* was shaped by co-existing microbial communities^53^ competing with *S. cerevisiae* in the enrichment media. The second part of the hypothesis, “but the environment selects,” relates to an outstanding question regarding whether *S. cerevisiae* has an ecological niche^52^. We show that two such substrate environments exist at a lineage level. TW4 was isolated only from fungal fruiting bodies and lichens, and a CHN-VI/VII.2 genetic component was present in many lineages and enriched in isolates recovered from the tree fruit substrate. Higher frequencies of admixed isolates observed in fruits may simply be a result of increased contacts with other lineages. Alternatively, fruits and organisms associated with those fruits such as frugivorous animals and vectors may represent niches that promote hybridization; for instance, sporulation has been suggested to be an adaptation that allows cells to survive in nutrient-depleted conditions such as insects’ intestines during experimental passaging^54^. Notably, the presence of CHN-VI/VII.2 genetic components in many natural lineages across the world, as well as in admixed isolates found in fruits, raises the possibility that the common ancestors dispersed from East Asia were from this lineage. In addition to abiotic factors, we speculate that such dispersal events of fruits may be aided by insects and human foraging.

We found that, unlike the general expectation in biogeographic studies that an island only contains a subset of genetic diversity from the mainland population, the genetic diversity of *S. cerevisiae* populations from Taiwan can be as diverse as those found in the Asia continent. The persistence of ancestral lineages may be a result of Taiwan being a high environmentally heterogeneous region^55,56^ and its prolonged bioclimatic stability^57^ than that of nearby eastern China. Alternatively, the geographic scale for distinguishing island and mainland populations and the importance of habitat diversity may differ between microorganisms^58^ and other macroorganisms, such as animals and plants. The biogeography of *S. cerevisiae* appears to be similar to that of its associated flora in East Asia. Disjunct distributions of plants between Taiwan and different parts of China are common^59^. The phylogeography of representative herbaceous and woody plants indicates that these representatives originated in mainland China, then migrated to Taiwan and the Ryukyu Archipelago during the Pleistocene as sea level fluctuations yielded recurring landbridges^60–62^. Interestingly, we note that the Pleistocene was also the period when severe tree species extinctions first took place across both the Americans^63^ and Europe^64^; this was followed by a rapid migration of *Quercus* that made it the dominant tree genus^64^, which may have played a role in the restricted *S. cerevisiae* lineages observed outside East Asia. A systematic sampling of *S. cerevisiae* in the mainland continent—especially regions containing flora records exhibiting a disjunct distribution like in Taiwan, e.g., the Himalaya-Hengduan mountains^62^, as well as plate boundaries—may help us better understand the biogeography of *S. cerevisiae*.

Our findings of rampant hybridization events between wild, wild with domesticated and domesticated lineages bring new perspectives to the ongoing debate over whether *S. cerevisiae* domestication happened once^2,65^ or multiple times^1,21^. By revealing frequent hybridizations between natural lineages, we show that isolates used in Asian and European fermentations may have been domesticated independently from the lineage CHN-VI/VII.2, and the single-domestication-event notion may be confounded by admixed isolates. Isolates from Asian fermentations were sister to the CHN-VI/VII.2 clade, and subsequent genetic differentiations of this group have led to independent lineages such as the North American oak group, or the Mediterranean oaks group which is sister to the European/Wine isolates (**Fig. 2a and 3a**). Isolates outside of East Asia likely bear genetic components of this group. This may result in the placement of these isolates in or close to this group in a phylogeny. Ongoing hybridizations also complicate the inference; for instance the Brazilian rum population is a result of hybridization between European/Wine and North American groups^21^. Efforts to identify signatures of domestication environments^65^ may also be challenging when admixture is detected between these lineages. Isolation and tracking the frequencies of these admixed isolates in nature could provide further insights into the conditions in which new lineages emerge.

To conclude, we combined deep sampling, metabarcoding, isolate collection and whole-genome resequencing to illuminate the pre-domestication phase of *Saccharomyces cerevisiae* at an unprecedented resolution. We reveal that multiple natural lineages of *S. cerevisiae* persist in Taiwan; the species is found everywhere but some genetically differentiated lineages prefer certain substrates. These observations help us to revisit our understanding of eukaryotic microorganism evolution—for instance, an alternating life cycle seems to be a convenient life history trait when genetically diverged partners are around. As more and more ecosystems—e.g., tropical cloud forests^66^—and biodiversity are lost, actions should be taken to conserve and reveal the ecology and evolution of not just *S. cerevisiae* but species with a proposed geographical origin. The availability and recurrent gene flow between these lineages also allow future experiments such as on hybrid fitness to be designed to resemble the subject’s natural scenarios rather than relying on domesticated strains.

## Methods

### Sampling and isolating *Saccharomyces cerevisiae*

From September 2016 to October 2020, we collected a total of 2,461 environmental samples from various substrates (bark *n*=340, twigs *n*=328, leaf *n*=528, litter *n*=320 and fruit *n*=78) surrounding 693 plant hosts (**Supplementary Table 2**). A total of 339 lichen samples, aliquots from six fermentation practices, and 68 from other sources (insect corpse *n*=43, fruiting body *n*=14, industrial strains *n*=5 and others where biomaterial was sampled only once *n*=6) were also collected. Collection time and GPS coordinate in gpx format of host plants was recorded on the day of collection. Leaves, flowers of host plants were photographed. Bioclimatic variables of sampling sites were retrieved from CHELSA^67^ database (v1.2) using recorded GPS coordinates. Digital terrain models (DTM) of sampling sites were retrieved from Taiwan’s Open Government Data website (https://data.gov.tw/dataset/35430). Environmental samples were collected using alcohol sterilized tweezers or spoons and stored in zip bags. Whenever possible during the sampling trips metadata such as the identity of the host plant, lichens and altitude were recorded. Samples were redistributed into 50ml falcon tubes and stored at room temperature. Each sample was divided into two proportions and immersed in two enrichment media: a liquid medium made up of either i) 3 g/L yeast extract, 3 g/L malt extract, 5 g/L peptone, 10 g/L sucrose, 7.6% EtOH, 1 mg/L chloramphenicol, and 0.1 % of 1-M HCl as used in ref^12^ or ii) YPD containing 10% dextrose and 5% ethanol adjusted to pH 5.3 as used in ref^68^. Samples were incubated at 30°C until signs of microbial growth and fermentation were detected, such as white sediment and effervescence. Sediments were then streaked onto YPD agar plates. Single colonies were picked out and incubated in potassium acetate medium 23°C for 7-10 days^69^. Single colonies with ascus-like (four spores) structures under microscope were picked out and streaked onto YPD agar plates. Sanger sequencing and gel electrophoresis of ITS1-5.8S-ITS2 region PCR amplified with ITS1F/ITS4 primer set were performed to identify the species of isolates^70,71^. Pilot sampling, modification and rationale during the course of sampling strategies are further provided in **Supplementary Info**. Sampling efforts were visualized using the R’s package ggplot2 (v.3.3.5) and annotated with metR (v.0.10.0; https://github.com/eliocamp/metR) and ggspatial (v.1.1.5; https://paleolimbot.github.io/ggspatial/). In order to determine ploidy levels for our isolates, we carried out flow cytometry analysis for the 105 Taiwanese isolates from this study using propidium iodide (PI) staining assay using previously established protocols^72^ (**Supplementary Info**).

### DNA extraction

Field-collected environmental samples can vary, so we preprocessed these samples and extracted their DNA differently (see **Supplementary Info** for details). For whole genome sequencing of *Saccharomyces cerevisiae*, isolates taken from frozen stocks were streaked out onto YPD plates and incubated in 30°C until colonies became visible. Single colonies were then incubated in 5ml YPD liquid medium at 30°C in a shaker at 200 rpm overnight. High molecular weight genomic DNA was extracted using protocol described in ref^73^. DNA quality was determined by Qubit readings, A260, A280, A260/280 ratios on Nanodrop and gel electrophoresis.

### Library construction and whole-genome sequencing

For Illumina sequencing, paired-end libraries were constructed using the Illumina Nextera or NEB Next Ultra DNA library preparation kit with the manufacturer’s protocol. The first 91 isolates were sequenced by Illumina HiSeq2500 and the remaining 30 were sequenced by Novaseq to produce 125- and 150-bp paired-end reads, respectively. Oxford Nanopore libraries were prepared using SQK-LSK109 with 12 isolates multiplexed by EXP-NBD104 and EXP-NBD114 barcoding kit (ver NBE_9065_v109_revV_14Aug2019) and sequenced by a R9.4.1 flow cell on a GridION instrument. A total of 24 isolates were run on two flow cells. Nanopore fast5 files were basecalled using Guppy (v4.0.11).

### Amplicon sequencing and analysis

Amplicon libraries were constructed as previously described^74^ from 89 environmental samples (18 bark, 18 twig, 18 leaf, 18 litter, 17 soil), three positive controls (*Saccharomyces cerevisiae* S288C, *S. paradoxus* YDG197 and lab isolate *Pseudocercospora fraxinii*); and DNA from two *Escherichia coli* as a template to confirm primer specificity towards only fungal species. The ITS3ngs (5- CANCGATGAAGAACGYRG-3’) and ITS4ngsUni (5’- CCTSCSCTTANTDATATGC-3’) primer pair^75^ was used. Two no template controls were included during the PCR step to confirm that amplicon generation was free of contaminating DNA. To determine the background amplicon noise from experimental pipeline, a sterile filter was treated and processed as one the field samples. Amplicons were normalized using the SequalPrepTM Normalization Plate Kit (ThermoFisher, ID: A1051001), then pooled and concentrated using AMPure XP (Beckman Coulter, ID: A63881). Finished DNA libraries were sequenced on the Illumina Miseq platform using 2×300 bp pair-end sequencing chemistry.

Raw sequencing reads containing the Illumina sequencing index were demultiplexed using *sabre* (v1.0; https://github.com/najoshi/sabre). Sequencing quality was determined using *FastQC* (v0.11.7; https://github.com/s-andrews/FastQC). Reads were quality filtered based on a Qscore >20 and 50 base pairs were trimmed from the 3’end end using usearch^76^ (v11.0.667). Filtered reads were processed following the UPARSE^77^ pipeline. In brief, paired reads were merged and dereplicated into unique sequences. Unique sequences were filtered using usearch default settings. Filtered sequences were denoised into zero-radius operational taxonomic unit (zOTUs) using the unoise2^78^ algorithm. The taxonomy of zOTUs was classified using the SINTAX^79^ algorithm (Edgar 2016) against the UNITE^80^ Fungal database (v8.2). Merged reads were assigned into zOTUs with 100% sequence identity and tabulated using the *usearch_global* function. Processed reads were analyzed in the R-Studio environment (v 1.2.5033). Sequencing data were analyzed with phyloseq^81^ (v1.34). Statistical significance was tested for using *kruskal.test* from the *stats* package in R (v4.0.2).

### Variant calling

To determine the evolutionary history of new Taiwanese isolates, we collected a total of 219 published genomes representing established *S. cerevisiae* industrial and natural populations: 102 isolates from the 1,011 genome project^1^ (31 Wine/European, 8 Mediterranean oak, 6 African beer, 6 African palm wine, 4 West African cocoa, 4 Malaysian bertam palm nectar, 6 North American oak, 6 Sake, 11 Asian fermentation, 1 CHN-I, 1 CHN-III, 4 CHN-IV, 1 CHN-V, 6 Mixed origin groups and 7 other isolates of Taiwanese origin), 93 isolates from the Chinese population^2^ (69 CHN-I to CHN-X isolates excluding those previously sequenced in the 1,011 genome project, 5 isolate from Mantou1, 6 Huaugjiu, 7 Baijiu and 6 Qingkejiu), 16 isolates from the Brazilian wild lineage^33^ and eight isolates from olive brine^34^. This combined with the 121 isolates from this study yielded a total of 340 individuals, 30% of which originated from industrial sources and 70% from the natural environment (**Supplementary Table 6**). Read quality was examined with FastQC (v.0.11.9; https://github.com/s-andrews/FastQC). Read quality and adaptor trimming was performed using Trimmomatic^82^ (v0.36; Pair end mode, ILLUMINACLIP;LEADING:20;TRAILING:20;SLIDINGWINDOW:4:20;MINLEN: 150). For the 340 samples, 64-95% of raw paired reads from the 340 samples were kept after trimming. Trimmed reads were each mapped to the S288C reference genome version R64-2-1 using Burrows-Wheeler Aligner^83^ (v 0.7.17-r1188) and the mapping rate was 91-99%. Duplicate reads were marked using gatk^84^ MarkDuplicates (v.4.1.9.0). Variants were first called in multi-sample manner and filtered using bcftools^85^ v1.8 (−d 1332; QUAL 30, MQ 30, AC >=2 and 50% missingness; genotype-filtered with minDP 3). 88% (1,150,658/1,306,082) of variants were retained. Second, variants were also called and filtered with freebayes^86^ (v. 1.3.2; minDP 3, QUAL 30, MQ 30, AC >=2 and 50% missingness, sites with 0.25<AB<0.75 and 0.9<MQM/MQMR<1.05 were retained). 56% of sites were retained based on these criteria (818,025/1,443,685). Finally, 808,864 intersecting variants discovered from both callers were used for further analysis. The functional effects of variants were annotated with SnpEff^87^ (v.4.3t).

### Assembly, annotation and ortholog identification

Nanopore reads of each isolate were assembled using Canu^88^ (v1.9). For isolates without long reads, Illumina paired-end reads were assembled using SPAdes^89^ (v. 3.14.1, options k-mer size 21, 33, 55, 77 and --careful). Consensus sequences of the assemblies were polished with four rounds of Racon^90^ (v1.4.11), one round of Medaka (v1.0.1) using nanopore raw reads and five rounds of Pilon using Illumina reads. The assemblies were further scaffolded using RagTag^91^ against the S288C genome reference. Annotations were then transferred using Liftoff^92^, with additional *de novo* annotations using Augustus^93^ on regions without any transferred annotations. The assembly metrics and description of the nanopore assemblies are shown in **Supplementary Table 8**.

### Phylogenomic analyses

After removing 43,695 invariant sites resulting from ambiguous nucleotide codes among all isolates, the remaining 765,169 variable sites were used to construct a phylogeny for the 340 isolates. The resulting best-fit model was indicated by BIC to be TVMe+R3 first with IQ-TREE. In addition, a maximum likelihood phylogeny was inferred using IQ-TREE with the TVMe+R3+ASC model and a 1,000 ultrafast bootstrap approximation^94,95^.

To infer the *Saccharomyces cerevisiae* lineage phylogeny, amino acid, nucleotide sequences and annotation of proteomes from the following species in the *Saccharomyces sensu stricto* clade were downloaded: *S. bayanus* FM1318 (Genbank accession GCA_001298625), *S. bayanus* CBS7001 (Genbank accession GCA_000166995.1), *S. kudriavzevii* NBRC 1802 and ZP 591 from ref^96^, *S. jurei* from ref^97^, *S. arboricolus* H6, *S. paradoxus* CBS432, N44, UFRJ50816, UWOPS91-917.1 and YPS138 from https://yjx1217.github.io/Yeast_PacBio_2016/data/. Orthology considering synteny information was inferred using PoFF^98^ (v. 6.0.27). The protein alignment was constructed for each of the 1,594 single copy ortholog groups using MAFFT^99^, then back-translated into a nucleotide sequence alignment using PAL2NAL^100^ (v.14). A maximum likelihood phylogeny was constructed with each of the 1,594 single copy ortholog protein alignments using RAxML-ng (v1.0.0, option --model LG+I+F+G4 --tree pars 10). The phylogeny and bootstrap support replicates were used together to infer a lineage phylogeny using ASTRAL-III^101^ (v5.6.3). A separate maximum likelihood phylogeny was built using RAxML-ng (v1.0.0) with the concatenated alignment of the single copy orthologs. This phylogeny and concatenated nucleotide alignment of the single copy orthologs were used as input for the MCMCtree method in the PAML^102^ package to estimate the divergence time among the *S. cerevisiae* lineages. The overall substitution rate was estimated from PAML^102^ based on the concatenated nucleotide alignment. The following molecular divergence estimates from ref^103^ were used to calibrate the phylogeny: *S. cerevisiae-S. paradoxus* 4–5.81 million years ago (Ma), *S. cerevisiae-S. mikatae* 6.97–9.47 Ma, *S. kudriavzevii-S. mikatae* 10.1–13 Ma, *S. arboricola-S. kudriavzevii* 11.7–14.8 Ma and *S. eubayanus-S. uvarum* 4.93–7.93 Ma.

### Diversity, population structure and demography estimates

For the population structure estimate, biallelic SNPs were kept and filtered based on linkage disequilibrium. Sites that are linked were filtered out using PLINK^104^ (v1.90b4), excluding pairs of loci with r^2^ > 0.5 (--indep-pairwise 50 10 0.5 --r2). The remaining 482,161 sites were used for ancestry estimation by ADMIXTURE^36^ using K=2 to K=30 with five-fold cross validation from five runs of different seed numbers. CV errors for each K value in five runs were compared to choose the representative number of clusters. Migration signals on the phylogeny were estimated with TreeMix using 1000 bootstrap for natural populations according to clusters in K=16. The numbers of migration edges were estimated, aided by the optM (v. 0.1.5) package in R (https://cran.r-project.org/web/packages/OptM) and presented in **Supplementary Info**. EEMS^40^ (v.6fa1aff commit) was run using distance matrix of Taiwanese and Asian wild isolates calculated from fastme^105^ (v. 2.1.5.1).

A consensus genome sequence containing variants for each isolate was generated from the SNPs matrix using bcftools^85^ consensus (v.R64-2-1) with the S288C reference genome sequence. Diversity estimates for 16 nuclear chromosomes and corresponding coding/noncoding regions were examined by VariScan^106^ with RunMode 11 (n<4) and 12 (n>=4), and custom python scripts. The recombination rate ρ=4N_e_r for each isolate was estimated by rhomap^107^ as part of the LDhat program (v2.2) with 10,000 iteration and samples taken every 100 iterations. Inbreeding coefficients F was determined for each isolate by PLINK^104^ (v.1.90b4; --ibc) on LD-trimmed SNP matrix. Together, using the relationship N_ρ_=ρ/(4r(1-F) and N_θ_=θ(1+F)/4μ (where SNV mutation rate was estimated as μ=2.82×10^−10^ from ref^108^), frequencies of sexual reproduction can be estimated as N_ρ_/ N_θ_ for each population. To estimate the ratio of nonsynonymous to synonymous substitution rates (*d_N_/d_S_*) for each gene, nucleotide sequence alignment and its translated protein sequence alignment were aligned using PAL2NAL^100^ (v.14) and d_*N*_/d_*S*_ estimated with the codeml program in PAML^102^. For the IBD analysis, geographical distance between isolates was measured using the sf package in R for Taiwanese isolates with GPS records. For Chinese isolates, since GPS records were not available, we used approximate coordinates for each sample site (personal communication with authors of ref^2^). To estimate the maximum geographical distance within the Chinese lineage, we chose sample sites that were the furthest apart. For instance, in CHN-V, the distance between Shanxi and Hainan was used. However, for lineages sampled from only one site (CHN-II, CHN-IX), the largest range of the site was used as the maximum distance within the lineage.

## Acknowledgement

We thank Cheng-Ruei Lee for the insightful comments on the ADMIXTURE analyses. We thank Mao-Ning Tuaumu for the helpful suggestions on how to deal with bioclimatic variable data. We thank Cheng-Ruei Lee, Jun-Yi Leu, Dang Liu and John Wang for commenting on earlier versions of the manuscript. We thank Jun-Yi Leu for the experimental advice and Bo-Fei Chen for helping with the initial sampling trips. We thank Tze-Fu Hsu, Yi-Hsiu Kuan, H. Thorsten Lumbsch, Matthew Nelsen, for collecting/providing some of the biomaterials. We are grateful to the National Center for High-Performance Computing for its computer time and for letting us use its facilities. I.J.T. was supported by the Ministry of Science and Technology, Taiwan under grant 110-2628-B-001-027 and Career Development Award AS-CDA-107-L01, Academia Sinica.

## Authors contribution

I.J.T. conceived and led the study. T.J.L, Y.C.L and W.A.L carried out the sampling and isolation of *Saccharomyces cerevisiae*. J.P.H, C.L.H and K.F.C helped with the sampling and identified the lichen and plant samples. T.J.L, W.A.L, Y.F.L and H.M.K conducted the experiments. Y.F.L carried out the amplicon analyses. H.H.L., H.M.K and I.J.T. performed the sequencing and assemblies of the *S. cerevisiae* genomes. T.J.L, Y.C.L, H.H.L and I.J.T. carried out the population genomic analyses.I.J.T. carried out the phylogenomics analyses and the divergence time estimation. T.J.L and I.J.T. wrote the manuscript with substantial input from J.P.H, K.F.C and G.L.

## Data availability

Raw data on the 121 *S. cerevisiae* isolates were deposited in the National Center for Biotechnology Information (accession no. PRJNA755173). The accession numbers of the isolates are shown in **Supplementary Table 6**. GPS coordinates of the isolates are available upon request.

