## Supplementary Info for "Deep sampling of ancestral genetic diversity reveals *Saccharomyces cerevisiae* pre-domestication life histories"

### Supplementary Text

#### Sampling strategies

In 2016, we began sampling by collecting barks from Fagaceae trees around Academia Sinica campus (Taipei City, Taiwan) using four different media that were previously reported to be successful in the selective enrichment of *Saccharomyces* yeast species[1-3]. It was not until July 2017, when we had our first successful isolation, that we started collecting samples from various substrates (including leaves, bark, litter and soil at the base of the tree) of amenity *Quercus glauca* trees near the National Taiwan University (NTU; Taipei City, Taiwan) campus. We noticed that different substrates yielded different levels of success, but the overall isolation success per tree host increased. Preliminary repeated sampling and isolation also showed that two out of four enrichment media (Sniegowski *et al*[4], Hyma and Fay's high sugar[5]) gave higher isolation success. We initially sequenced 23 isolates on the same trees and found that these isolates were primarily clones of each other, though genetically differentiated isolates could still be identified (**Supplementary Table 10**). To ensure effective sampling efforts and avoid redundancies in sequencing, we decided to i) sample various substrates in a given tree host, ii) used two enrichment media and iii) sequence only one isolate if multiple isolates were recovered from same substrate from a tree site. Following this decision, we began to expand the sampling sites to look for natural isolates.

We first made a trip around Taiwan in December 2017 each collecting samples from two to five trees from different regions of Taiwan. These trees were mostly but not entirely belonged to the *Fagaceae* family. When an isolate was successfully recovered from a site, we would revisit that region to sample adjacent trees at least tens of meters apart, if the environment permitted. The whole cycle of round trip to new sites, isolation and revisiting was repeated several times until October 2020, when we had recovered what we believed the be enough isolates covering enough sites to yield statistically significant results. Species identification was initially carried out by multiplex PCR to detect *Saccharomyces cerevisiae* species-specific amplicons as described by ref[6], but was subsequently modified to add step 6-8 in ref[7] detailed in Methods. For species identification, single colonies were picked out and lysed in QuickExtract DNA solution (epicentre). The ITS1F (5-CTTGGTCATTTAGAGGAAGTAA -3') and ITS4 (5'-

TCCTCCGCTTATTGATATGC-3') primer pair were used. The PCR cocktail consisted of 1 µl Colony lysate, 1 ul each of the primers (5 µmol), 12.5 µl KAPA Taq Ready Mix PCR Kit (KK1024, Kapa Biosystems, USA), and 9.5 µl double-distilled water. The following thermocycling conditions were used: an initial 3 min at 95°C, followed by 30 cycles at 95°C for 30s, 52°C for 30s,72°C for 1min, and a final cycle of 5 min at 72°C, cooling at 16°C.

**Determining ploidy level of *S. cerevisiae* isolates**

Frozen yeast stocks were first streaked out on YPD plates and kept in 30℃ until colonies became visible. For each isolate, a single colony was picked out and grown in liquid YPD overnight at 30℃ with agitation (200rpm). Cultures were diluted in YPD to obtain $OD_{600}$ = 0.2, and incubated under the same condition for an additional 5-6 hours until $OD_{600}$ reached around 1. Each cell culture was diluted again with YPD to achieve $OD_{600}$ = 0.6, and 500µl of the liquid culture was transferred to a microcentrifuge tube. Cells were harvested with a benchtop centrifuge at 500 *x g* for 3 minutes at room temperature, then pellets were subsequently washed with 500µl sterilized dd$H_2O$. Cell pellets were collected again with centrifugation, resuspended in 1ml cold 70% ethanol and left to fix at 4℃ overnight. Cells were collected with centrifugation and resuspended in 50µl sodium citrate (50mM) with vortexing at max speed for 10 seconds, centrifuged again and resuspended in 200 µl sodium citrate (50mM) with a final concentration of 0.5mg/ml RNaseA (prepared in 40% glycerol). RNaseA treatment continued for 2 hours at 37℃. Cells were transferred into dark brown microcentrifuge tubes for subsequent staining. A final concentration of 25 µg/ml of propidium iodide was added to each sample, then samples were incubated at 37℃ overnight in the dark. Samples were vortexed at max speed for 10 seconds, then 25 µg/ml propidium iodide (in 50 mM sodium citrate) was added to create a 1:40 dilution of cells. Each diluted sample was then passed through a sterilized 30 µm pre-separation filter to remove any remaining large cell clumps. Cell cycles were recorded on a Beckman Coulter CytoFLEX S Cell Analyzer with a 610 nm filter. More than 20,000 events were recorded for each isolate, and standard samples with known DNA content were used as baseline to compare YL610-A(area) values at 2C and 4C. Ratios of mean YL610-A ranged between 1.78-2.24 for diploid isolates and 2.93 for one triploid isolate found in our collection.

**Collection of additional geographical data**

The last-level administrative divisions in Taiwan were retrieved from the GADM database (https://gadm.org/) and coordinates were grepped from Google Maps with R package RgoogleMaps[8] (v1.4.5.3).

**DNA extraction from environmental samples**

*Bark* 1.0 g of each bark sample was powdered using pestle and mortar with liquid nitrogen. 10 mL of lysis buffer (100mM Tris.Cl pH8, 1.4M NaCl, 2.0% w/v CTAB, 20mM EDTA and 1.0% w/v PVP)[9] was added to the powdered samples, which were then incubated for 1 hr at 65°C. Nucleic acids were extracted with 10mL of chloroform:isoamyl alcohol (24:1) mixed by inversion using a benchtop rotator at 30 rpm for 10min. Lysate mixture was centrifugated at 10,000 *x rcf* for 30 min at 25°C. The aqueous phase was transferred to a new tube, and nucleic acid precipitated with an equal volume of isopropanol. Precipitated nucleic acid was centrifugated at 10,000 *x rcf* for 10 min at 25°C. Supernatant was decanted, and the pellet air-dried for 5 min at room temperature. DNA was resuspended in 150 µL ddH$_2$O. Bark DNA was cleaned further using an equal volume of AMPure XP beads (Beckman Coulter, ID: A63881), per the manufacturer's protocol.

*Leaf, twig and litter* Due to variations in the field, collection sample size and surface were varied among the different tree families. Sample weights were not standardized to minimize handling and contamination. For each leaf, twig and litter sample, the quantity processed were weighed and transferred into a sterile 500 mL polypropylene centrifugation bottle (Beckman Coulter; Cat. 361691) using sterile tweezers. Samples were suspended in 250 mL of 1X PBS pH 7.4 with 0.1% Tween 20 and placed in a sonicating water bath (DELTA ULTRASONIC CO. LTD, Make: DELTA, Model: DC400) at a 40 kHz frequency for 20 min at 25°C as recommended by ref[10]. Sonicated samples were transferred onto a horizontal shaker (Double Eagle Enterprise Co., Ltd, Make: TKS, Model: OSI 500) set to 120 rpm for 1 hour at 25°C. Large debris was removed by passing the suspension through a 0.25 mm sterile mesh. Flow-through was centrifugated at 10,000 x *rcf* for 1 hour at room temperature. The supernatant was filtrated using a 0.22µm PES membrane filtration cup (Jet Bio-Filtration Co., ID: FPE214250). Pellets were resuspended using 1 X PBS with 0.1% Tween 20 and added to the filtration cup towards the end of filtration to minimise blockage. Filter membranes were excised using a scalpel, and total nucleic acid was extracted using a DNeasy PowerWater kit (QIAGEN; ID: 14900), per the manufacturer's instructions.

*Soil.* Soil samples were sieved through 2mm sterilized stainless steel mesh to remove visible rocks, insects, and plant materials. Sieved soils were homogenized by mixing using a sterile spatula. Total nucleic acid was extracted from 0.3g of processed soil samples using a DNeasy PowerSoil kit (QIANGEN, Cat. 12888) with minor modifications to the manufacturer's protocol. The incubation period after the addition of Solutions C2 and C3 was extended to one hour.

**TreeMix analyses**

We inferred the relationship between *S. cerevisiae* lineages using TreeMix[11] (v.1.13). Lineages were defined by different ADMIXTURE genetic compositions at different K values (16 or 29). At ADMIXTURE K=16, the following criteria were used: i) isolates with >97.5% genetic ancestry from one single group was designated to that group, ii) CHN-VIII with genetic component from CHN-VI/VII.2 and Wine/European was designated as a group, iii) TW3 with CHN-VI/VII.2 and CHN_X/Malaysian component was designated as a group, iv) TW6 with genetic component from African beer, Wine/European and Qingkejiu/Sake was designated as a group, v) five isolates recovered from steamed buns (Mantou) containing genetic component from African beer and Qingkejiu/Sake, and vi) PD38A was designated as a group. At ADMIXTURE K=29, the following criteria were used: i) isolates with >97.5% genetic ancestry from one single group was designated to that group, ii) TW6 isolates were designated as a group, and iii) PD38A was designated as a group. To ensure the independence of the sites, PLINK[12] (v1.90b6.20) was first run with options --indep-pairwise 50 10 0.5 and the block size in TreeMix was set to 100. TreeMix was run from one to 10 migration events with TW1/CHN-IX was used as the outgroup of the phylogeny. Five independent replicates were run on each edge to assess the consistency of the inferences. When the number of migration events in the phylogeny was increased from one to 10, the variance explained in the model increased by up to 99.6% at K=16 (**Supplementary Figure 7a**). We inferred seven migration edges as the most likely model (**Figure 3a**), as it explained 99.3% of the variance calculated using an *ad hoc* statistic based on change in the log likelihood between models of incrementing edges (**Supplementary Figure 7b**). Based on the K=29 designated grouping, we inferred eight migration edges to be the most likely model (**Supplementary Figure 9**) explaining 98.9% of the variance (**Supplementary Figure 10**).

### Supplementary Figures

**Supplementary Figure 1 – Sampling of 53 trees near Nanshan Village, Yilan County, Taiwan.** A total of three sampling trips were made to this location. Point denotes each sampled tree, colors denote whether and when the isolate was recovered, and shape denotes the sampling time. A total of 18 trees were sampled in both the second and third trip; these were annotated with numbers.

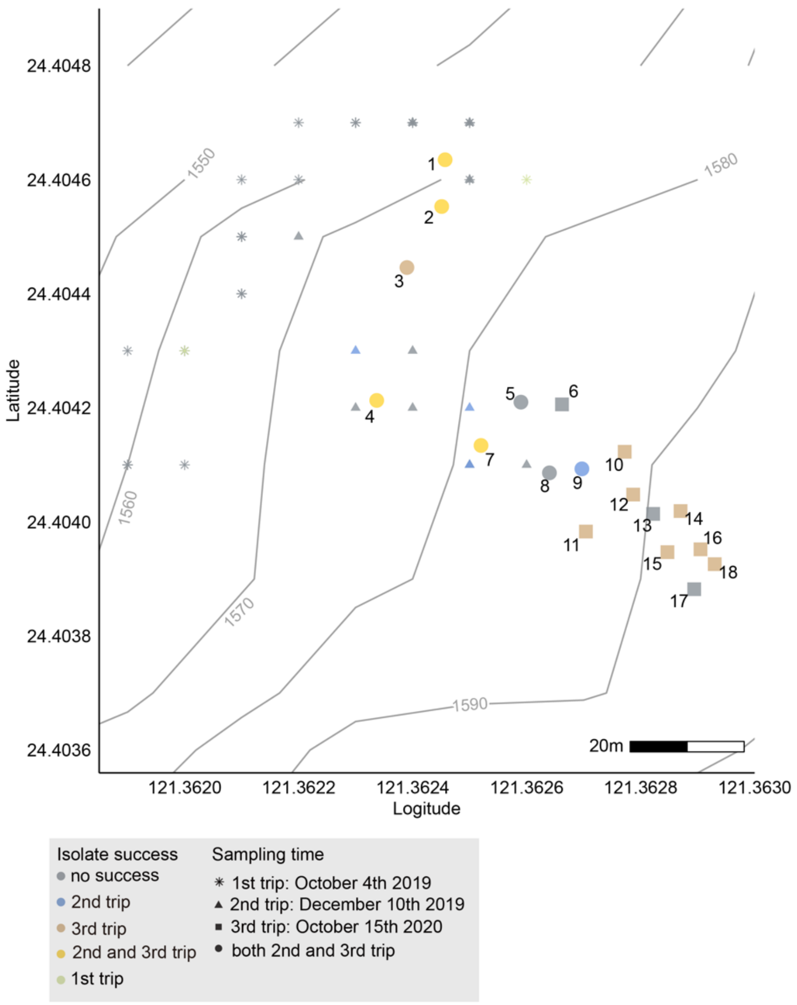

**Supplementary Figure 2 – *Saccharomyces cerevisiae* relative abundance amongst tree families.** The percentage abundance of *S. cerevisiae* relative to the total fungal abundance per sample was calculated. The relative abundance of *S. cerevisiae* amongst tree families are shown in a boxplot. Samples collected from different parts of the trees were denoted by the different shapes and colours. No statistical significances was detected between tree families (Wilcoxon rank sum test, P=1.0).

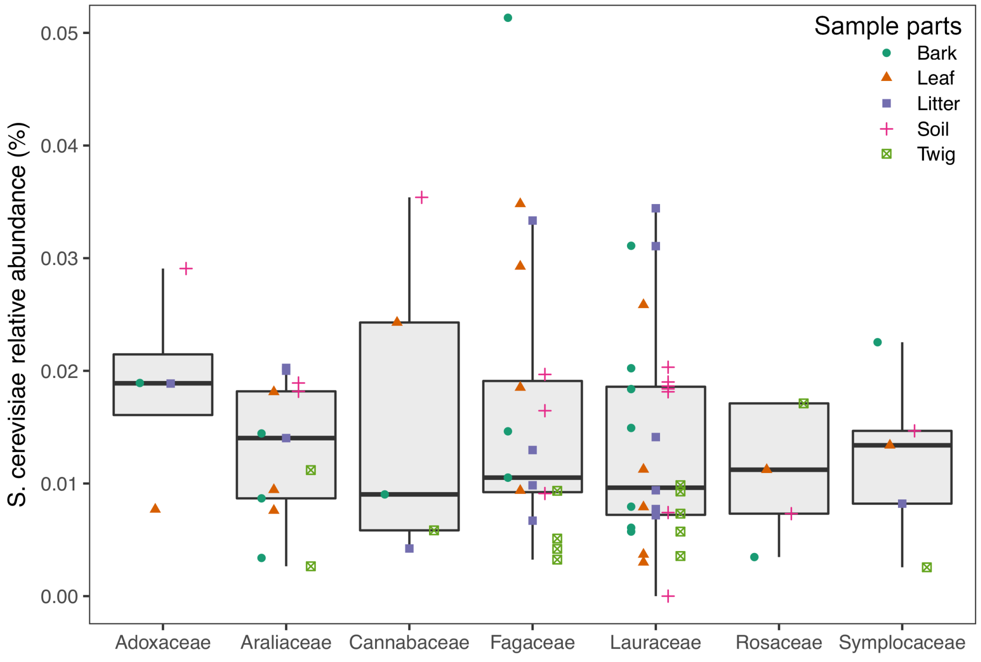

**Supplementary Figure 3 – Fluorescence histogram of diploid and triploid isolates.** Fluorescence histogram of yeast cells stained with propidium iodide of diploid (brick) and triploid (blue) isolates. Around 20000 events were sampled and filtered with successive gates: non-debris(P1), singlet cells(P2), and cells in G1/G2 phase.

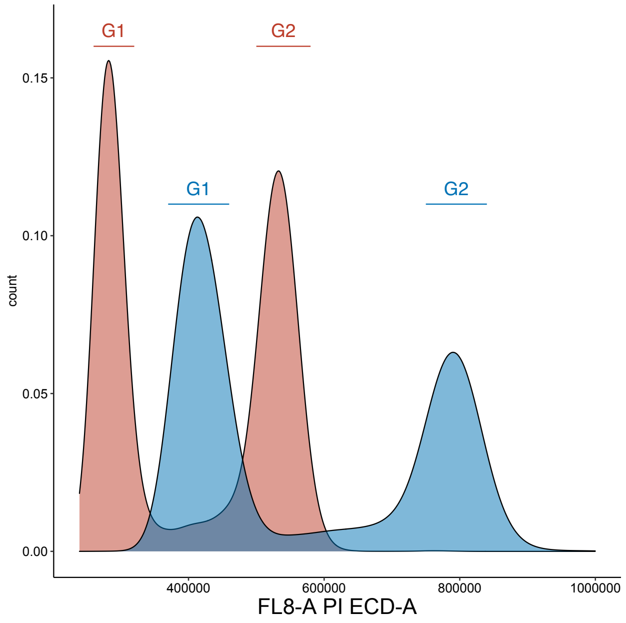


| Sample | P1 Events | P2 Events | G1 Events | G2 Events | G1:Median PI-A | G2:Median PI-A |
|---|---|---|---|---|---|---|
| 2N | 20000 | 18333 | 9227 | 8506 | 276418.8 | 525596.9 |
| 3N | 20012 | 18507 | 9943 | 7122 | 415769.8 | 792873.3 |

**Supplementary Figure 4 – Cross validation (CV) error estimates from ADMIXTURE output for *S. cerevisiae* 340 isolates.** Different point colors denote the five independent ADMIXTURE[13] runs from K=2 to K=30 with a line connecting the CV error of the first run.

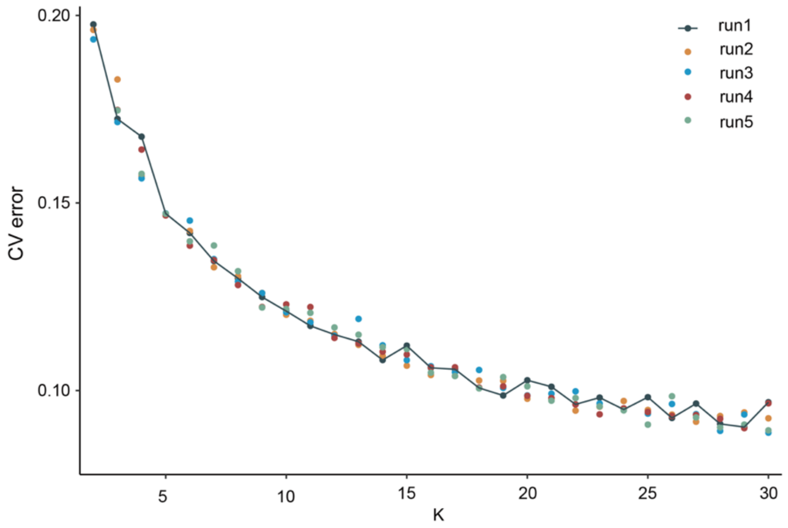

**Supplementary Figure 5 – Percentage of CHN-VI/VII.2a genetic component of ADMIXTURE K=29 analysis in *S. cerevisiae* natural isolates.** These isolates were categorized by substrate source and geographical origin (C and T denote Chinese and Taiwanese origin, respectively). N denote number of isolates.

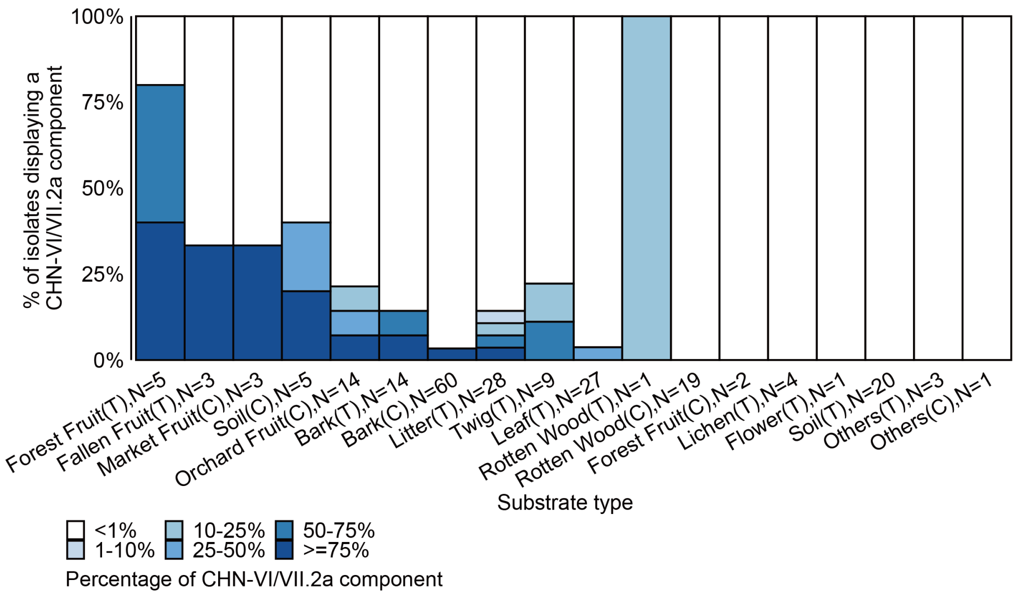


**Supplementary Figure 6 – Estimated effective migration surfaces (EEMS)[14] for wild *S. cerevisiae* isolates.** Migration rates (*m*) are color-contoured on a $\log_{10}$ scale. A blue color of 1 indicates an effective migration rate 10-fold faster than average. Green circle denotes locations where CHN-VI/VII.2 isolates was present.

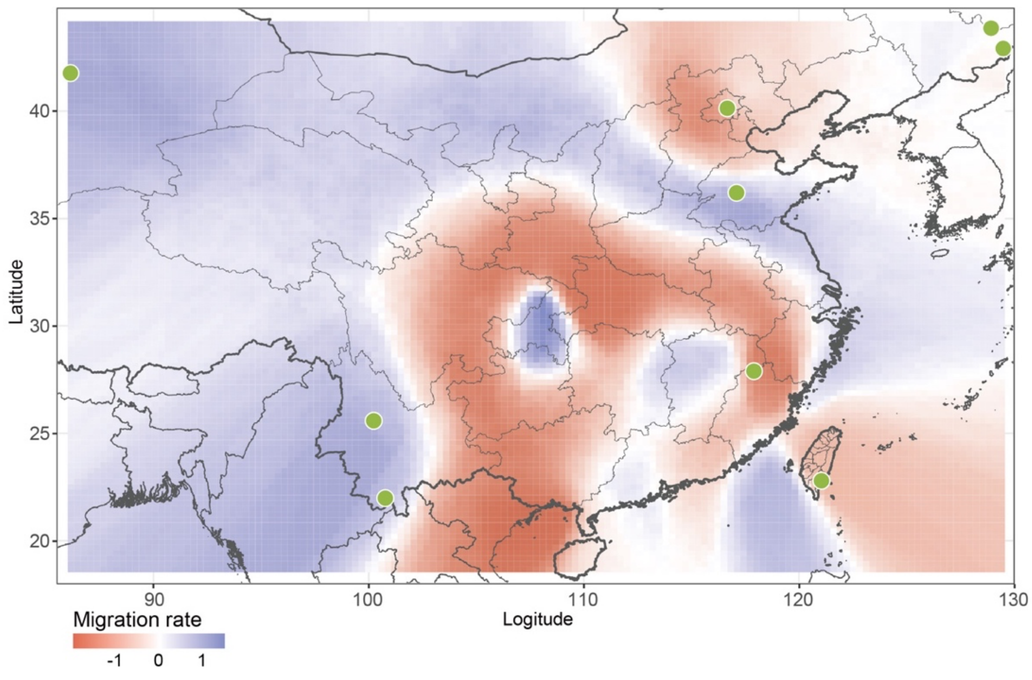

**Supplementary Figure 7 – Selecting the most likely model of ADMIXTURE K=16 based on the OptM R package.** a. Distribution of log likelihood and variances explained by models with 0–10 edges. Standard deviations generated by independent TreeMix runs of varying k values (1, 5, 10, 50, 500, 1000). b. Distribution of deltaM– an ad hoc statistic based on the second order rate of change–in the log likelihood with standard deviation considered. We inferred seven edges (>99% variance explained plus the second highest deltaM) to be the most likely model based on ADMIXTURE K=16.

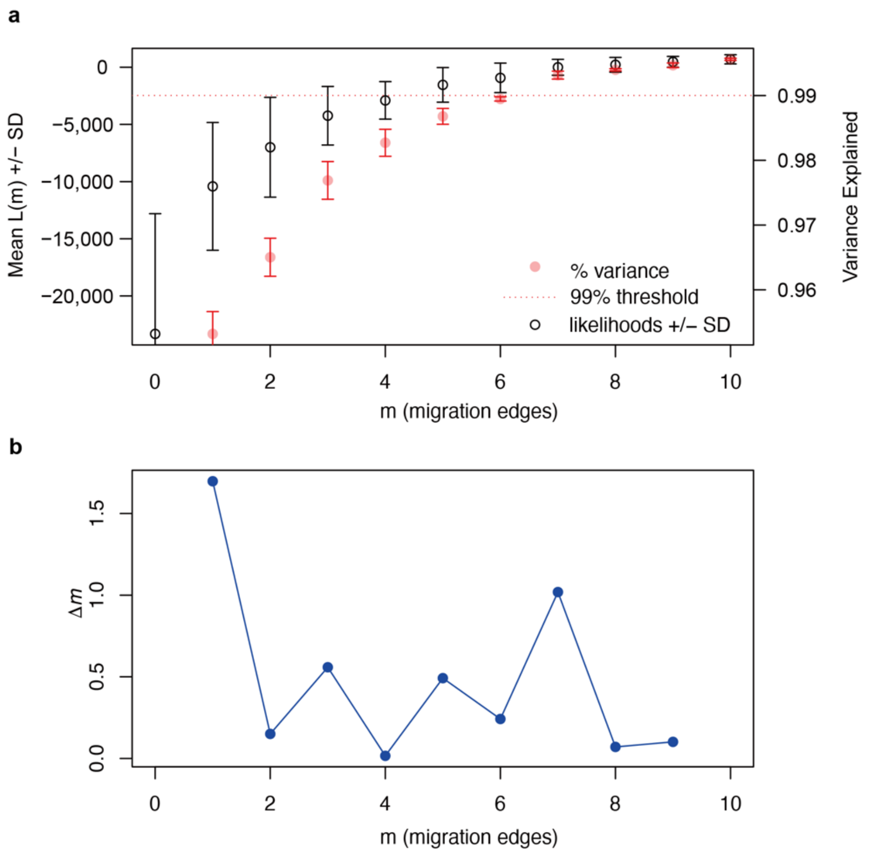

**Supplementary Figure 8 – Pairwise divergence over 10kb non-overlapping windows across 16 nuclear chromosomes among isolate PD35A (TW2 lineage), PD36A (TW4 lineage) and PD38A (hybrid of TW2 and TW4 lineage).** Divergence K was calculated using VariScan[15] (v2.0.3., RunMode=21).

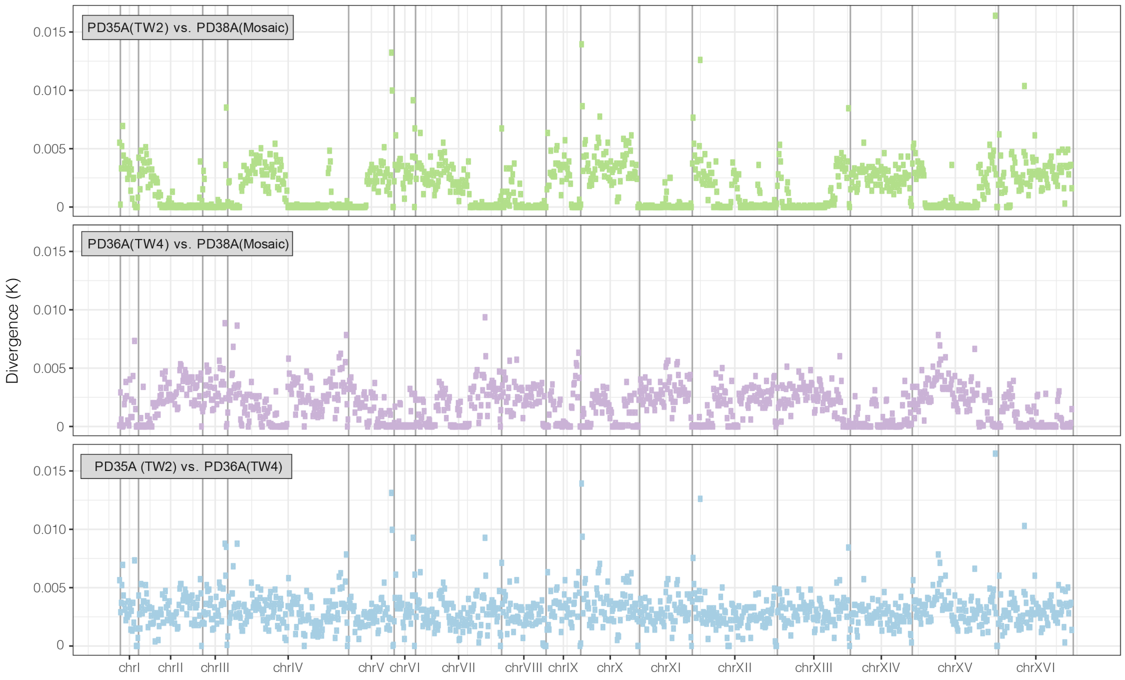

**Supplementary Figure 9 – The estimated relationships among the *S. cerevisiae* lineages with eight migration edges based on ADMIXTURE K=29 results.**

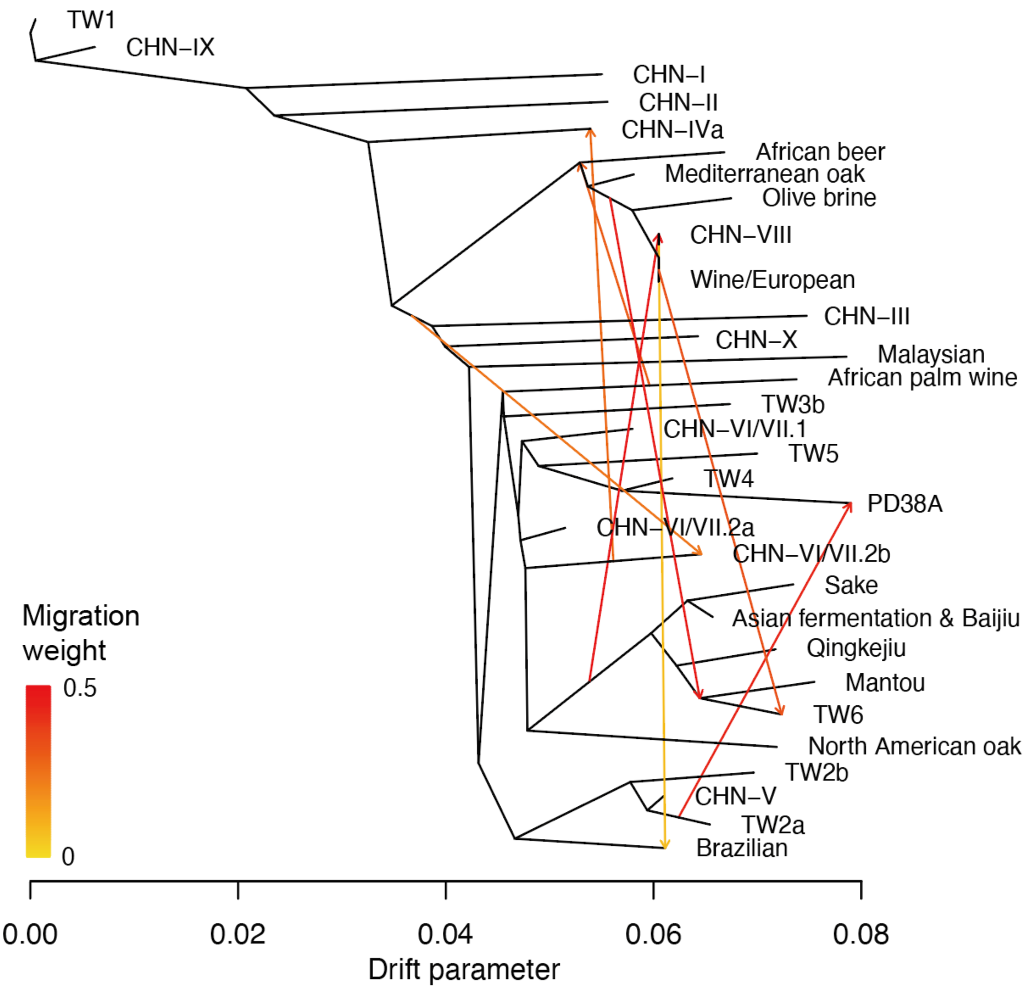

**Supplementary Figure 10 – Selecting the most likely model of ADMIXTURE K=29 based on the OptM R package.** a. Distribution of log likelihood and variances explained by models with 0–10 edges. Standard deviations generated by independent TreeMix runs of varying k values (1, 5, 10, 50, 500, 1000). b. Distribution of deltaM– an ad hoc statistic based on the second order rate of change–in the log likelihood with standard deviation considered. We inferred seven edges (~90% variance explained plus the third highest deltaM) to be the most likely model based on ADMIXTURE K=29.

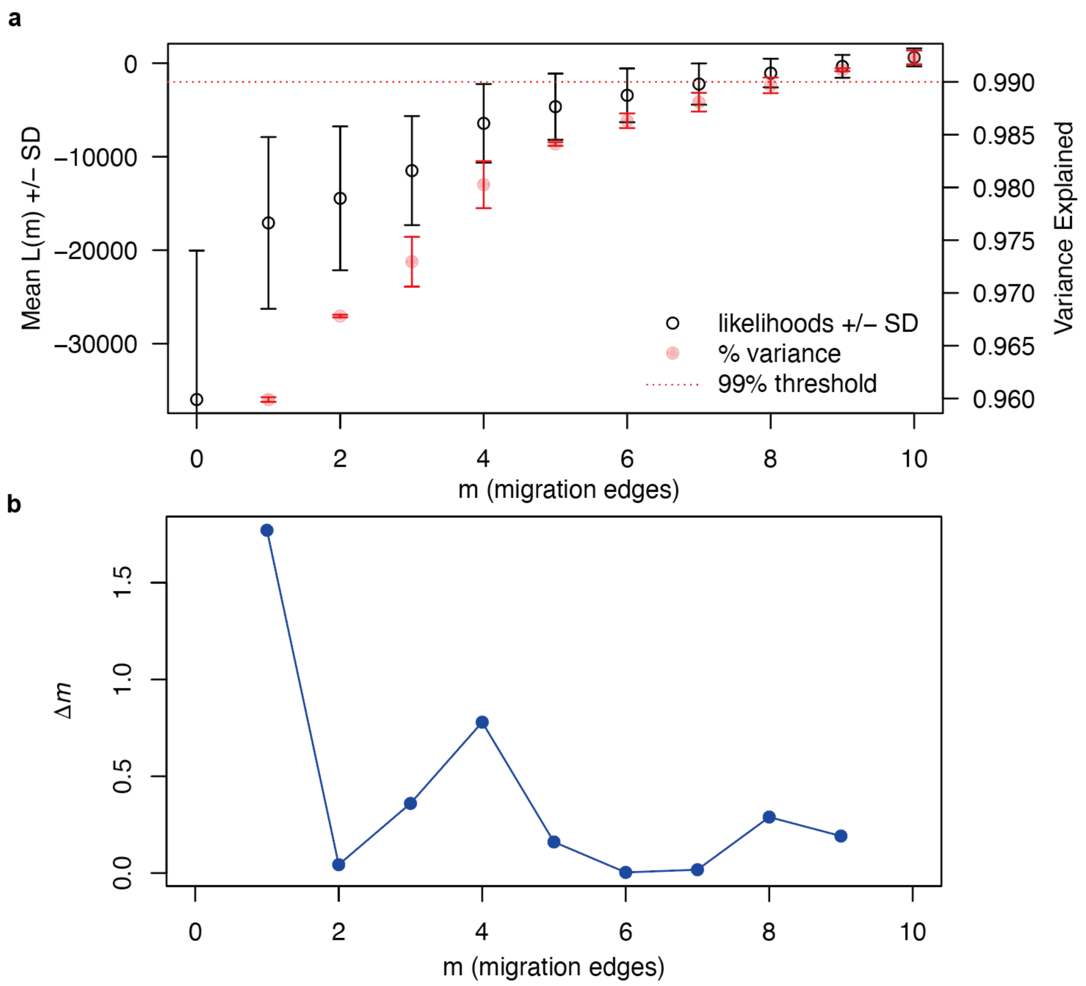

**Supplementary Figure 11 – *S. cerevisiae* lineage phylogeny inferred from a coalescence of 1,594 single-copy orthogroup gene phylogenies using ASTRAL[16].** Blue points denote selected nodes with unambiguous time divergence estimates shown on **Figure 3b**.

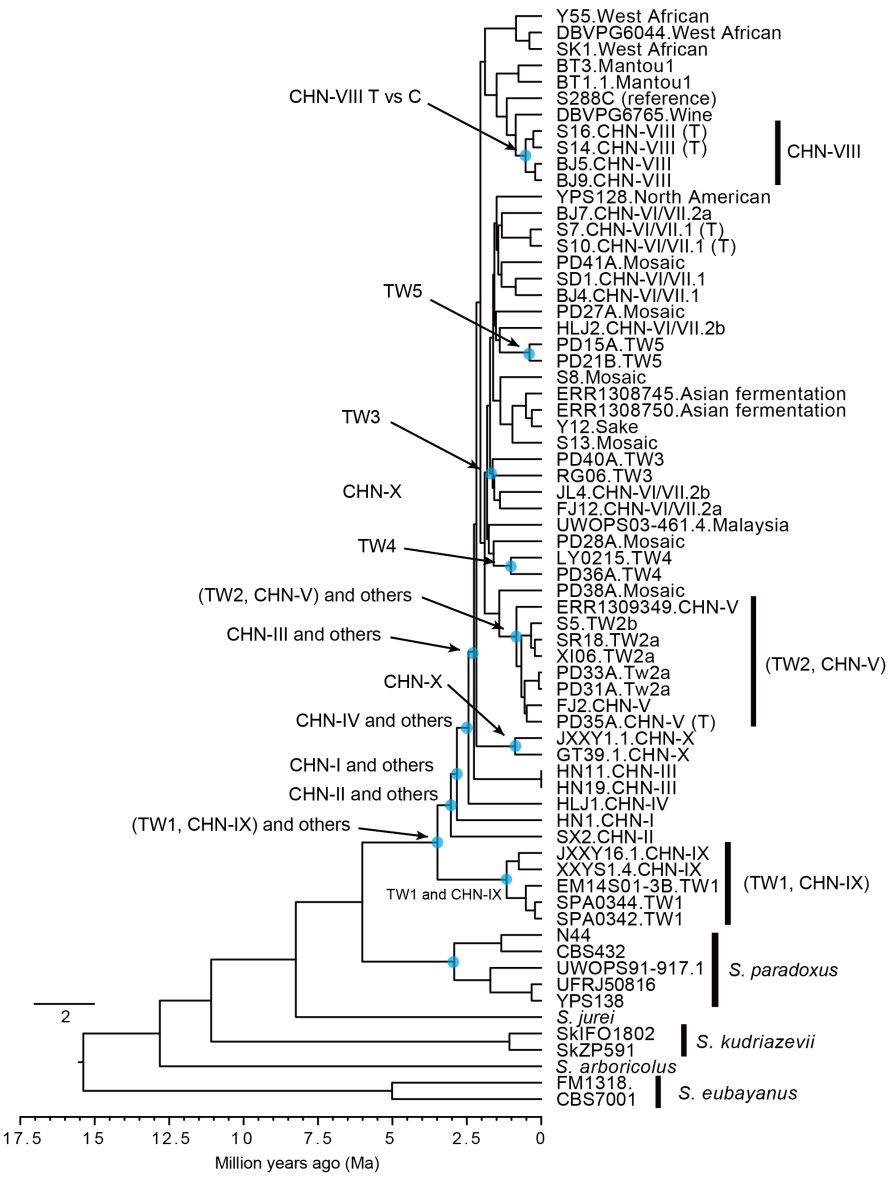

**Supplementary Figure 12 – Sampling of 105 trees near Puli, Nantou County, Taiwan.** Filled circles denote sampled trees without *S. cerevisiae* isolated. Some points overlap completely because of close proximity on the map. Arrow indicate the location of the tree that has five isolates from different lineages recovered.

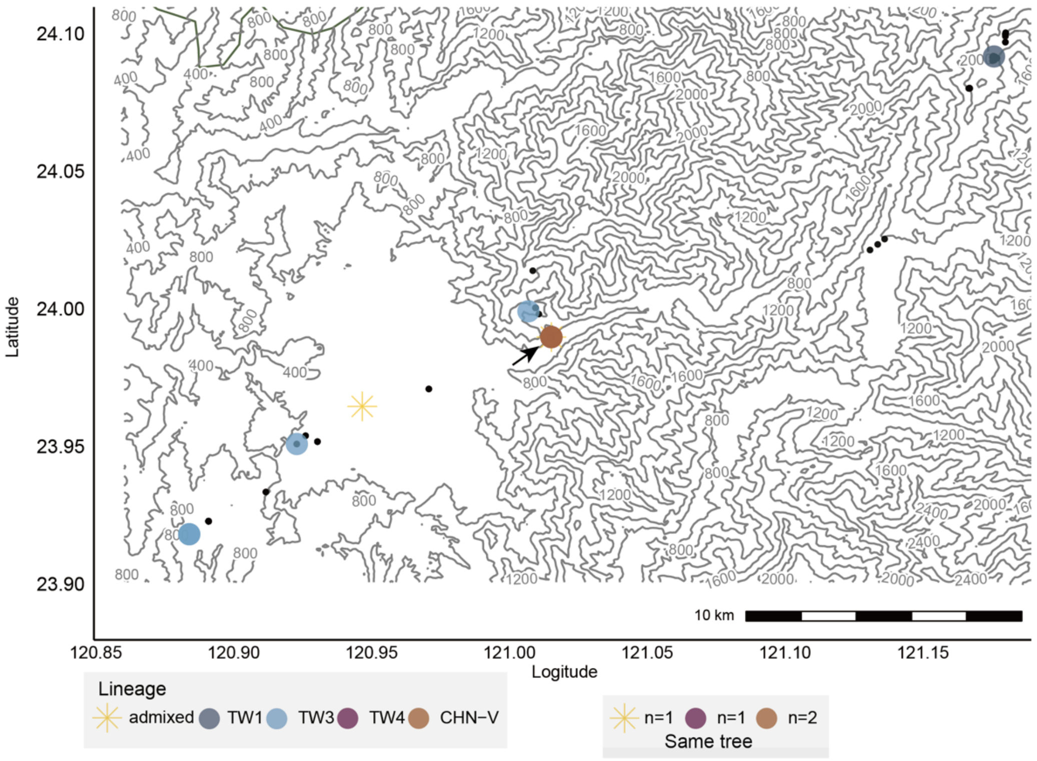


**Supplementary Figure 13 – Mantel correlation r for each geographic distance class in Fushan Botanical Garden.** Filled squares are statistically significant ($p<0.05$).

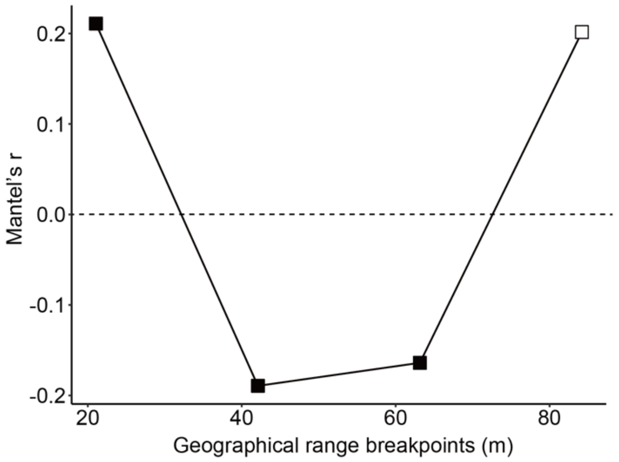

**Supplementary Figure 14 – Admixture proportion of the four admixed isolates in Fushan.** Genetic makeup of the four admixed isolates found in Fushan Botanical Garden estimated by ADMIXTURE at K=29.

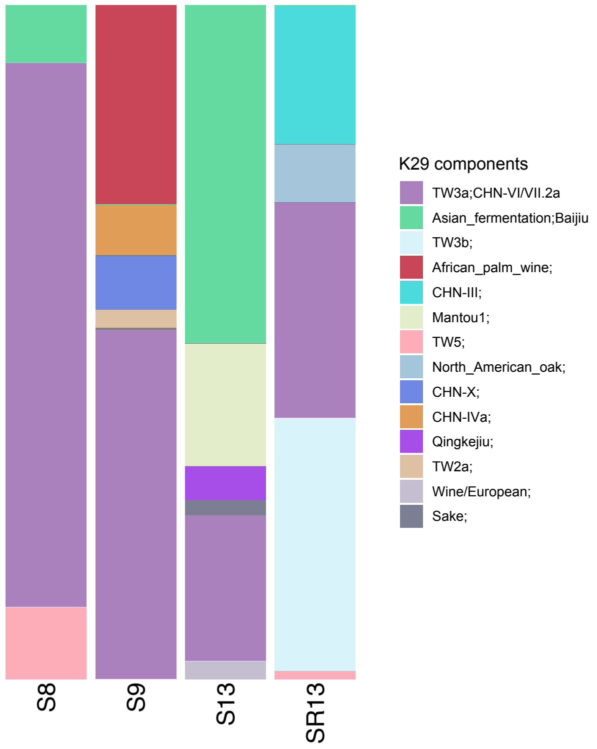


**Supplementary Figure 15 – Genetic diversities $\theta_\pi$ in noncoding region among isolates within a sampling area, and overall diversity for all Taiwanese isolates.** Dashed horizontal line indicates the overall diversity among natural Chinese isolates, which spanned over 3,500 km.

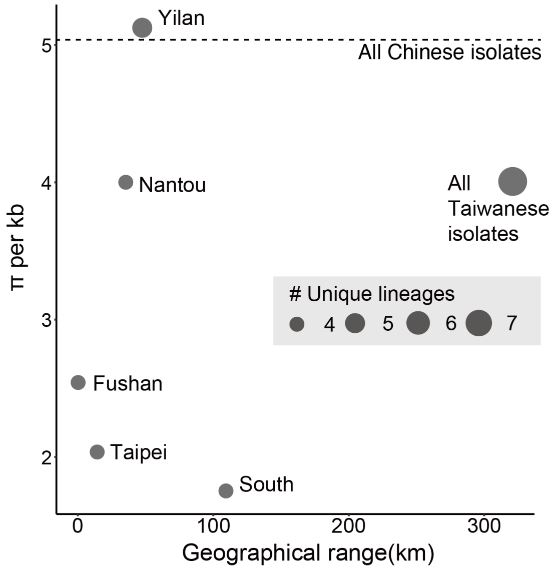

**Supplementary Figure 16 – Pearson's r between genetics and geographical distance across natural Taiwanese lineages.**

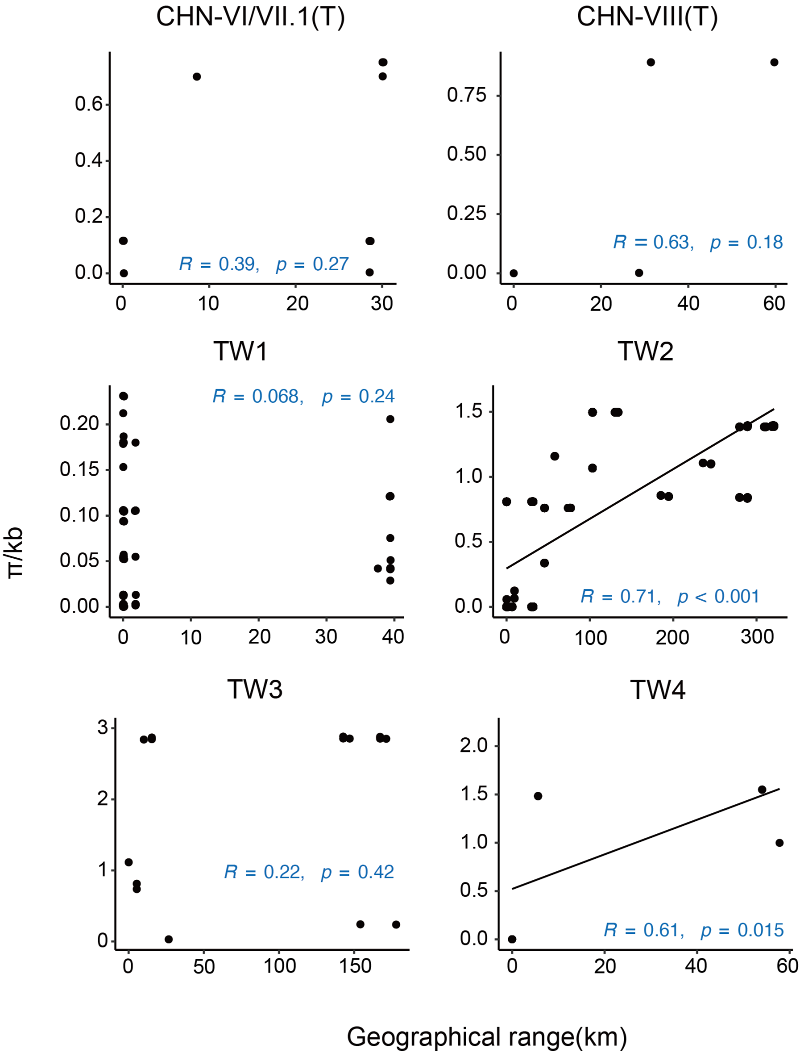

**Supplementary Figure 17 – Pairwise sequence diversity in relation to increasing pairwise geographical distance** Pairwise sequence diversity between natural Taiwanese isolates in incremental geographical distance categories.

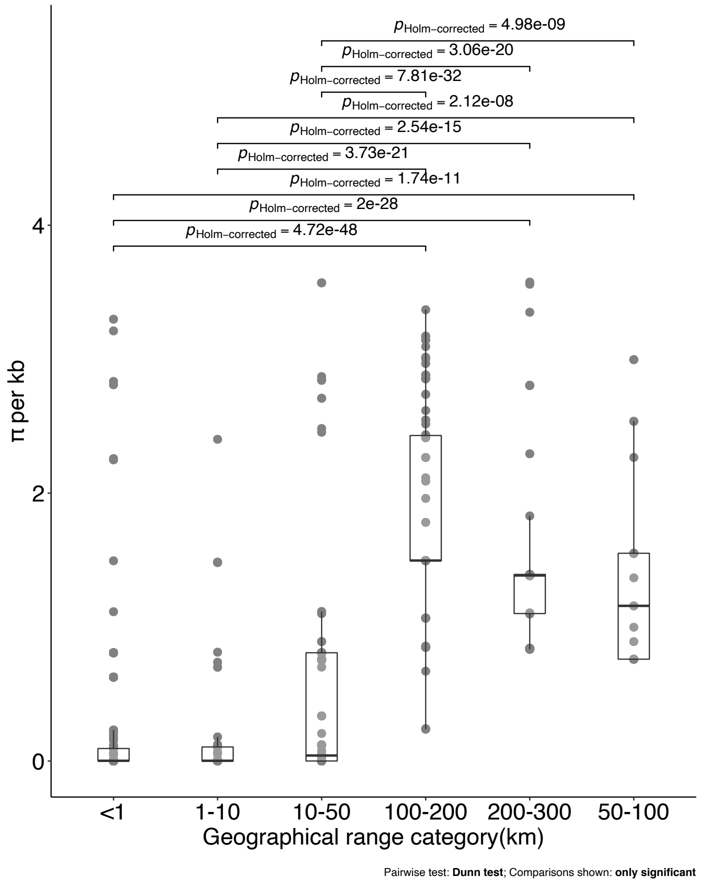

**Supplementary Figure 18 – Number of genes with *dN/dS*>1 in each of the five groups.** Number of genes with *dN/dS*>1 in isolate pairs from monophyletic TW-CHN groups. Single filled circles represent genes unique to only one group, connected circles show genes that are present in at least two groups.

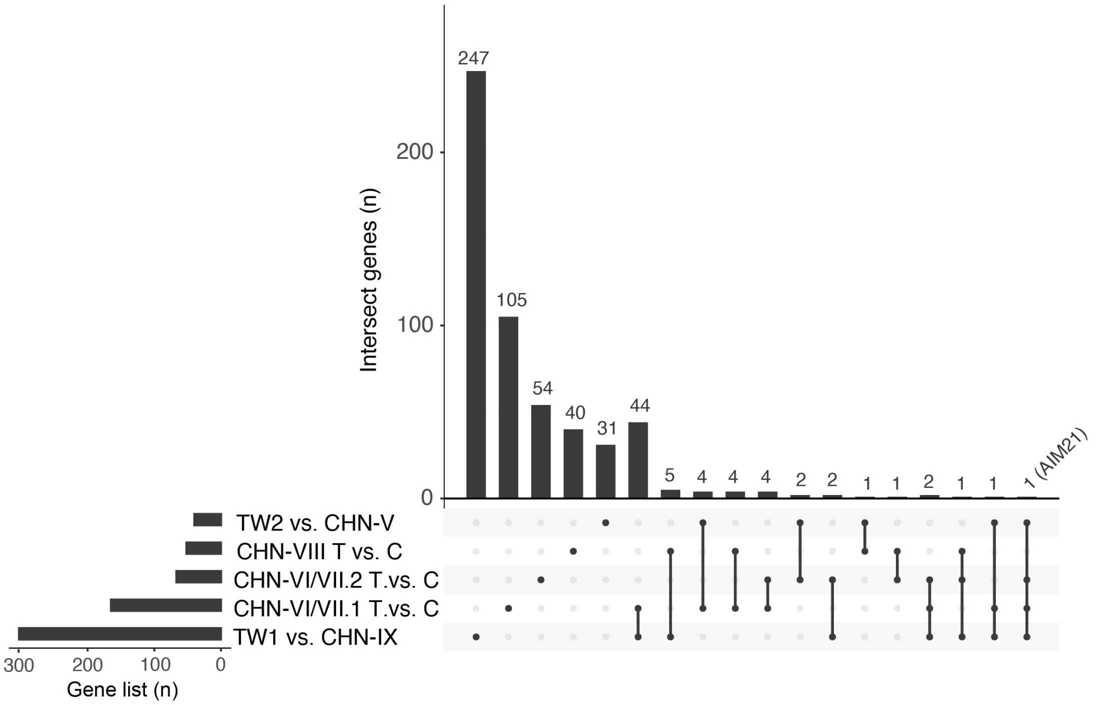

**Supplementary Figure 19 – Frequency of alleles across lineages when CHN/TW pairwise comparisons were $F_{ST}$ = 1.** Allele frequency distribution in lineages for SNP with $F_{ST}$=1 in any of the five comparisons (TW1:CHN-IX; TW2:CHN-V; CHN-VIII(T):CHN-VIII(C); CHN-VI/VII.1(T): CHN-VI/VII.1(C))

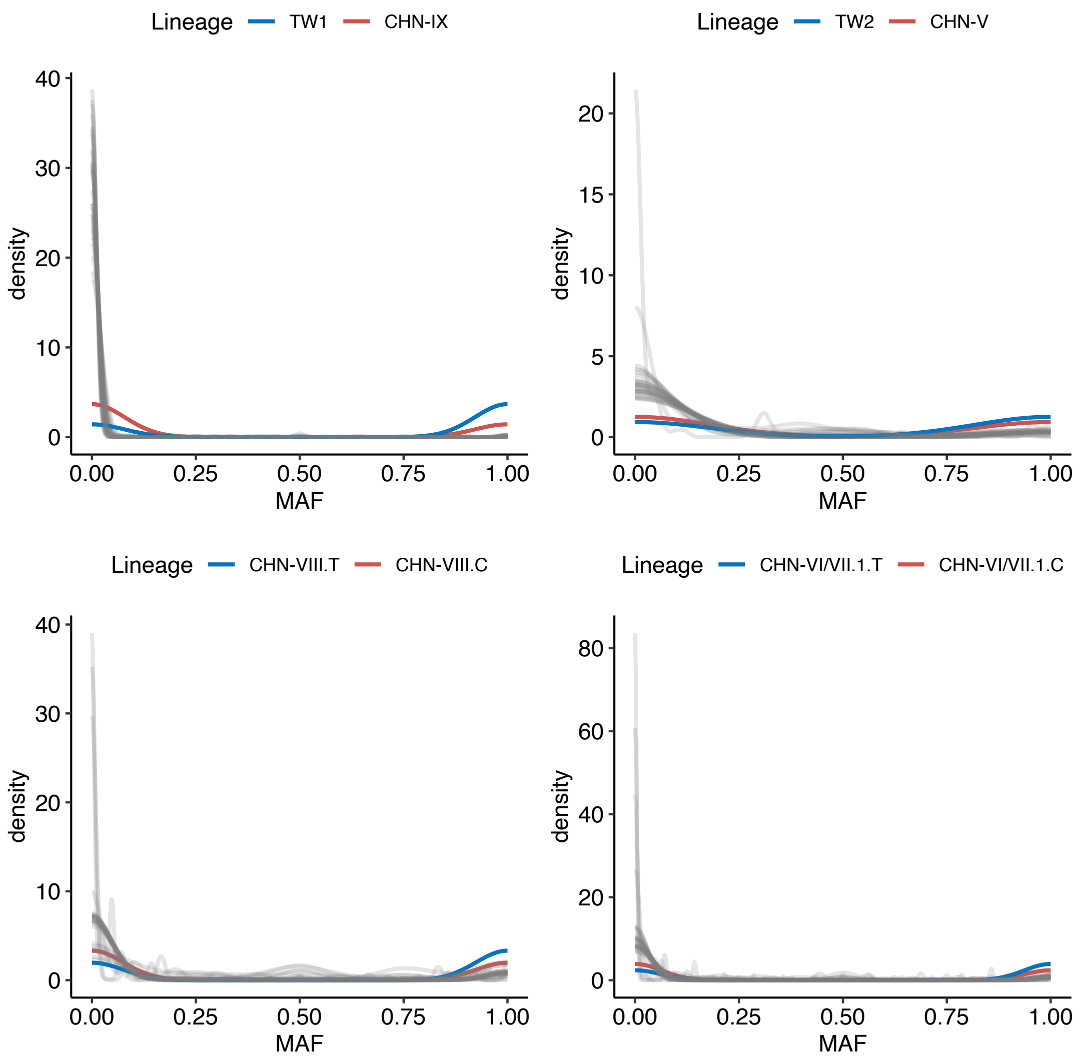

### Supplementary Tables

All tables are saved in a merged Excel xlsx file

**Supplementary Table 1 Descriptions of samples and yeast isolates collected in this study**

**Supplementary Table 2 Detailed information on sampled plant hosts and associated isolation success rates.**

**Supplementary Table 3 Isolation success rates in different types of samples, and in different months of the year from plant samples.**

**Supplementary Table 4 Isolation success rate for repeatedly sampled trees between two years**

**Supplementary Table 5 No significant difference in bioclimatic variables between regions with or without yeast isolates**

**Supplementary Table 6 Details on the 340 isolates used in this study including previously published genomes.**

**Supplementary Table 7 The 340 isolates, their phylogenetic groups and the lineages corresponding to their top three major components estimated by ADMIXTURE.**

**Supplementary Table 8 List of isolates with assemblies produced from nanopore reads**

**Supplementary Table 9 Range estimate for divergence time for selected nodes on the phylogeny in Supplementary Figure 10.**

**Supplementary Table 10 Regional genetic diversity estimates by VariScan[15] and maximum geographical range for natural lineages**

**Supplementary Table 11. Genetic diversity among isolates recovered within the same tree host**

**Supplementary Table 12 Ranges of effective mutational and recombinational population size, rate of sexual reproduction and number of asexual generations per sexual generation in natural lineages**

**Supplementary Table 13 List of genes with dN/dS>1 in pairwise comparison of Taiwanese and Chinese isolate of shared lineages.**

**Supplementary Table 14 Number and effect of SNPs with FST=1 in each lineage**
